# RIG-I RNA agonist activates immunostimulatory macrophages to enhance checkpoint immunotherapy for glioblastoma

**DOI:** 10.64898/2026.01.07.698153

**Authors:** Han Xu, Sungwoon Lee, Felipe Leser, Olga Fedorova, Peiwen Lu, Eric Song, Mehdi Touat, Anne Eichmann, Akiko Iwasaki, Anna Marie Pyle, Jean-Leon Thomas

**Affiliations:** Department of Neurology, Yale University School of Medicine, New Haven, CT, USA; Cardiovascular Research Center, Department of Internal Medicine, Yale University School of Medicine, New Haven, CT, USA; Department of Molecular, Cellular and Developmental Biology, Yale University School of Medicine, New Haven, CT, USA; Department of Immunobiology, Yale University School of Medicine, New Haven, CT, USA; Department of Ophthalmology and Visual Science, Yale University School of Medicine, New Haven, CT, USA; Sorbonne Université, AP-HP, Institut du Cerveau - Paris Brain Institute (ICM), Inserm U1127, CNRS UMR 7225, Service de Neuro-oncologie, Hôpital Pitié-Salpêtrière, Paris, France; Department of Cellular and Molecular Physiology, Yale University School of Medicine, New Haven, CT, USA; Paris Cardiovascular Research Center, Inserm U970, Université Paris, Paris, France; Center for Infection and Immunity, Yale University School of Medicine, New Haven, CT, USA; Howard Hughes Medical Institute, Chevy Chase, MD, USA; Department of Chemistry, Yale University, New Haven, CT 06511, USA

**Keywords:** glioblastoma, tumor-associated macrophages, pattern recognition receptor, RIG-I, stem-loop RNA, SLR14

## Abstract

Glioblastoma (GBM), the most frequent and aggressive primary brain tumor, remains refractory to all current therapies including surgical resection, chemotherapy, radiotherapy and immunotherapy. Immunosuppressive mechanisms in the GBM tumor microenvironment contribute to the lack of anti-tumor adaptive immunity. We found that a subset of tumor associated macrophages (TAMs) can be repolarized into an anti-tumor phenotype via agonist stimulation of the retinoic acid-inducible gene I (*RIGI*), a cytosolic double-stranded RNA pattern recognition receptor (PRR). In silico analysis of adult GBM datasets available in the public domain revealed that *RIGI* expression by a subset of activated TAMs positively correlated with patient survival. Studies in syngeneic mouse models of GBM showed that intratumoral delivery of stem-loop RNA 14 (SLR14), a RIG-I agonist, improved the efficacy of chemotherapy, radiotherapy and immunotherapy treatments, beyond the effects of other nuclei acid sensor agonists. We found that *RIGI*^+^ macrophages are the main drivers of SLR14 effect, combining activation of TAMs and priming of functional cytotoxic CD8^+^ T lymphocytes and NK cells. The anti-GBM effect of SLR14 is opening a significant new avenue for adult GBM treatment.

---

Glioblastoma multiforme (GBM) is the most aggressive form of primary brain tumor, with an incidence of approximately 4 cases per 100,000 people in the United States^1^. Despite advances in surgery, radiotherapy and chemotherapy, the median survival of GBM patients remains only of 12 months^2^. Cancer immunotherapies—including immune checkpoint inhibitors, tumor vaccines, and chimeric antigen receptor T-cell (CAR-T) therapy—have transformed cancer treatment and achieved clinical success in several tumor types^3,4^. Unfortunately, the immunotherapies tested thus far have failed to improve clinical outcomes in unselected cohorts of patients with GBM^5^. A key reason for the limited efficacy of immunotherapies in GBM is the strongly immunosuppressive tumor microenvironment^6^. GBM is marked by poor infiltration of effector T cells^7^, the accumulation of immunosuppressive populations such as tumor-associated macrophages (TAMs)^8^ and regulatory T cells^9^, and elevated expression of inhibitory molecules, including PD-1 and CTLA-4^10,11^. TAM populations are particularly abundant in GBM, where they can constitute up to 40% of the tumor mass and are major drivers of tumor-associated immunosuppression. Thus, there is increasing interest in understanding how the innate immune system shapes the GBM microenvironment. In tumors, immune activation is triggered by the sensing of cellular stress, and damage-associated molecular patterns (DAMPs) coexist with strong immunosuppressive and tissue-repair programs^12,13^. Pattern recognition receptors (PRRs), such as RIG-I-like receptors (RLRs), Toll-like receptors (TLRs), and cGAS–STING, are key detectors of DAMPs capable of initiating rapid protective responses^14,15^,and their activation enhances antitumor immunity by promoting myeloid activation and T-cell recruitment^16^. Among these strategies, the nucleic-acid-sensors TLR3/7/8 and cGAS have attracted interest in GBM, as preclinical^17–19^ and early clinical^20^ findings imply that stimulating innate immune recognition could help overcome the robust immunosuppressive tumor microenvironment. However, the potential of RIG-I signaling to the anti-tumor response remains largely unexplored, especially in the context of GBM^21–23^.

Here, we integrated human and mouse scRNA-seq data and identified a novel *RIGI*^+^ TAM subpopulation that expresses IFN-I related genes and derives from GBM infiltrated-monocytes. Preclinical studies using syngeneic mouse models of GBM further showed that the RIG-I agonist SLR14 exerts strong antitumor effects and improves the efficacy of combination treatments in GBM.

## Characterization of a subset of *RIGI*-expressing tumor suppressive macrophages in human GBMs

We previously developed a series of short stem-loop RNAs (SLR10 and SLR14)^24^ which potently induce type I interferon (IFNI) responses through RIG-I activation in vivo and drive robust antitumor effects in melanoma models^25^. To determine whether RIG-I is a potential target for anti-GBM immunotherapy, we first investigated *RIGI* expression in human GBM, using RNA sequencing (RNA-seq) databases from the Cancer Genome Atlas (TCGA). *RIGI* expression was increased in a subset of human cancer types, with GBM displaying a strong and statistically significant increase compared with adjacent normal brain tissue (Fig. 1a and Extended Data Fig. 1a). We next characterized the cell type-specific expression of *RIGI* from single cell RNA-seq (scRNA-seq) data derived from newly diagnosed grade IV GBMs (GSE182109, n = 8)^26^. Unbiased clustering and lineage annotation based on canonical marker gene expression allowed us to identify 75,499 single cells which segregated into thirteen major lineages spanning tumor, endothelial, stromal, and immune compartments (Fig. 1b and Extended Data Fig. 1b). *RIGI* expression was predominantly enriched in macrophages (Fig. 1c, d). Among the five clusters of tumor-associated macrophages (TAMs) identified from all GBM patients, the *RIGI*^+^ cluster exhibited prominent expression of IFN-stimulated genes (*IFIT1*) and IFN-response genes (*CXCL10* and *ISG15*). The *RIGI*^+^ cluster was clearly distinct from the cluster of MRC1^+^ macrophages that expresses non-canonical myeloid marker genes (*FOLR2*, *LYVE1*, and *SELENOP*) and corresponds to perivascular resident^27,28^ and tumor protective macrophages (Fig. 1e-f). Other clusters of TAMs expressed genes encoding for lipid metabolism (*APOE*) and immunosuppressive (*SPP1*)^29^, or inflammatory (*IL1B*) responses^13^. The high expression of *RIGI*-related genes (*RIGI*, *IFIT1*, *IFIT3*, *CXCL10*, and *ISG15*) was specifically associated with the tumor suppressive M1 TAM signature in GBMs, compared with other clusters (Extended Data Fig. 1c). Using analysis of spatial transcriptomic data derived from GBM samples (n = 6)^30^, we found that *RIGI*^+^ TAMs clusters tended to localize at the periphery of the hypoxic core within the tumor, within the angiogenesis-immune hub of the tumor microenvironment (Fig. 1g, h and Extended Data Fig. 1d, 3). Importantly, we could correlate the high expression of *RIGI*-related genes with a significant survival benefit for GBM patients (Fig. 1i). In contrast, elevated expression of *SPP1*- or *IL1B*-associated gene signatures correlated with poorer clinical outcomes (Extended Data Fig. 1f). Overall, these data justify further translational studies that are aimed at targeting *RIGI*^+^ TAMs for promoting macrophage polarization and anti-tumor response against GBM.

**Fig. 1.**
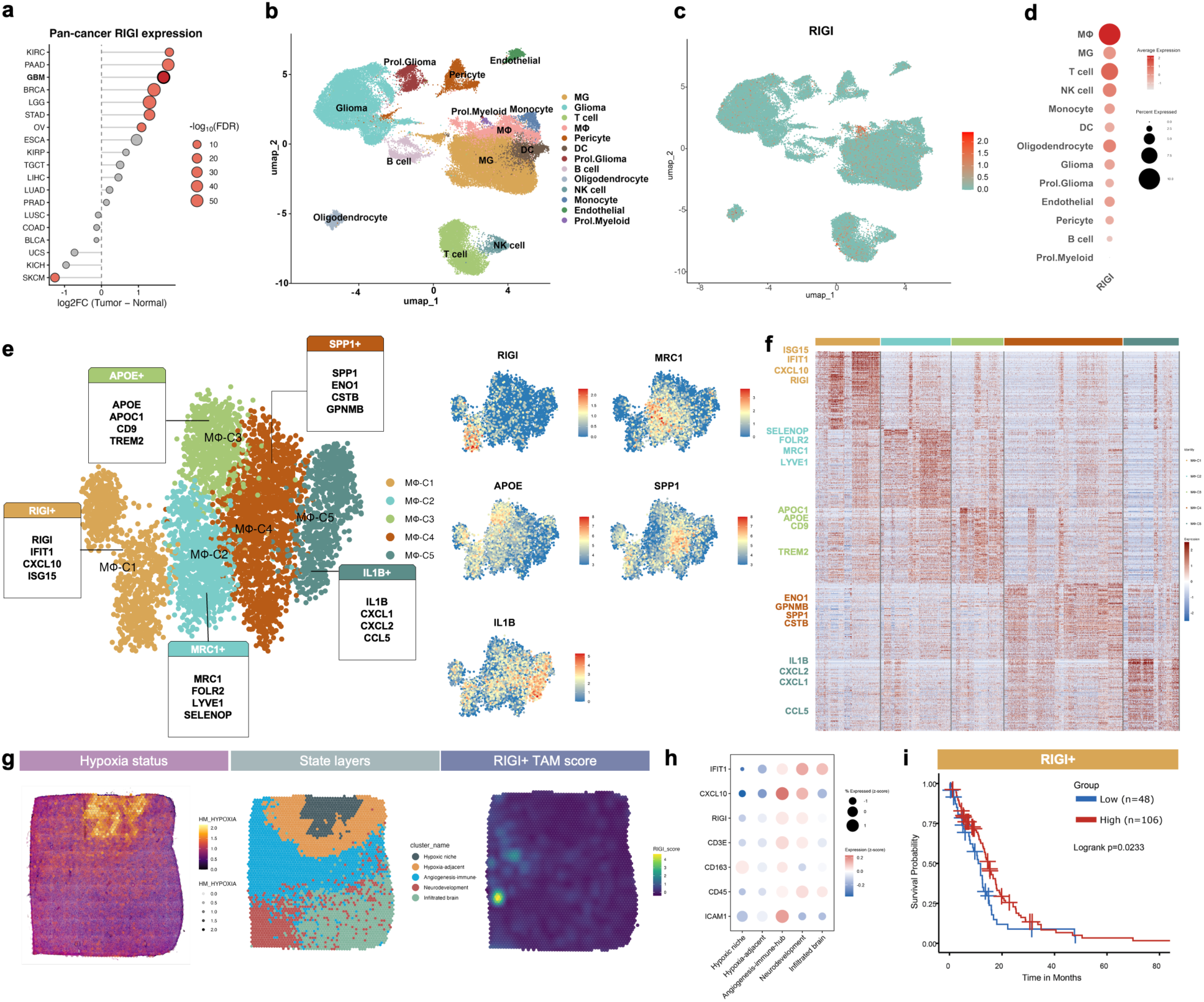
*RIGI* expression characterizes tumor protective innate immune cells and positively correlates with survival of patients with GBM. **a,** *RIGI* expression across various human cancers and their origin tissue, only tumors with significant change in *RIGI* expression are highlighted in red. **b**, Uniform manifold approximation and projection (UMAP) of scRNA-seq data from newly diagnosed GBM patients (n = 8, GSE182109). **c**, UMAP visualization of *RIGI* gene expression. **d**, Dot plots of scaled *RIGI* transcript expression in the indicated cell populations. **e**, UMAP of scRNA-seq data (GSE182109) from macrophages (left) and marker gene expression (right). **f**, Heat map of scaled expression of the top 50 marker genes for each macrophage clusters. **g**, Surface plots showing the transcriptional programs of the hypoxia area (left), the state layer (middle), and the *RIGI*-expressing TAM score (*RIGI*, *CXCL10*, *IFIT1*) (right) at spatial resolution using public hGBM spatial transcriptomic datasets (case UKF275 from the Dryad database). **h**, Dot plot showing *RIGI*-expressing TAM-related gene enriched in the angiogenesis-immune-hub (case UKF275 from the Dryad database). **i**, Survival probability of patients with GBM (from TCGA data) stratified with respect to the RIGI^+^ TAM gene signature.

## Conserved phenotype of *RIG-I*-expressing TAMs between human and mouse GBM

We next performed scRNA-seq analyses of GBM tumors collected from syngeneic GL261 GBM-bearing mice and age-matched controls to assess whether mouse GBMs also expressed RIG-I in TAMs. Twelve major cell lineages were identified, with a marked infiltration of immune populations, including macrophages, monocytes, dendritic cells (DCs), natural killer (NK) cells, and T cells (Extended Data Fig. 2a). Gene ontology (GO) analysis further revealed broad activation of both innate and adaptive immune response pathways in tumor-bearing brains (Extended Data Fig. 2b). Consistent with human GBM data, *Rigi* expression was predominantly enriched in TAMs (Extended Data Fig. 2c, d). We found that macrophage clusters were conserved between human and mouse GBMs, including *Rigi*⁺ TAMs, *Mrc1*⁺ TAMs, *Apoe1*⁺ TAMs, *Spp1*⁺ TAMs, and *Il1b*⁺ TAMs, which displayed transcriptional profiles like those of human TAMs (Extended Data Fig. 2e, f). *Rigi*⁺, *Spp1*⁺ and *Il1b*⁺ TAMs were significantly expanded after tumor inoculation, suggesting that GBM growth can induce activation of *Rigi*-expressing tumor suppressive macrophages (Extended Data Fig. 2g). In addition to macrophages, monocytes also showed a high *Rigi* expression in mouse GBM (Extended Data Fig. 2d), leading us to investigate a possible lineage relationship between *Rigi*-expressing TAMs and monocytes. Monocytes were characterized as Ly6c^hi^ cells (Extended Data Fig. 2h) and the inferred trajectory analysis suggested that *Rigi*^+^ TAMs, *Spp1*⁺ TAMs, and *Il1b*⁺ TAMs were more likely derived from monocytes than perivascular *Mrc1*⁺ TAMs^31^ and *Apoe1*⁺ TAMs (Extended Data Fig. 2i). Together, these findings identify *Rigi*^+^ TAMs as a conserved, monocyte-derived population that is markedly expanded in the microenvironment during GBM growth.

## Anti-GBM effects of intratumoral delivery of RIG-I agonist SLR14

The conserved presence of RIG-I⁺ TAMs in the microenvironment of human and mouse GBMs and the association between *RIGI*⁺ TAMs and improved outcomes in human GBM prompted us to assess the effect of RIG-I activation on survival in a mouse syngeneic model of GBM. We hypothesized that intratumoral delivery of a RIG-I agonist would promote the expansion of *RIGI*⁺/IFN-I secreting-anti-tumor TAMs. SLR14 is a known RIG-I agonist that induces IFN-I secretion by TAMs, suppresses peripheral tumor growth *in vivo*, and improves survival, particularly when combined with immunotherapies like anti-PD1 antibodies^25,32,33^. We first used the C57BL/6 syngeneic GL261 glioma mouse model to evaluate the antitumor efficacy of SLR14 against brain tumors in an immunotherapy-responsive setting (Fig. 2a). To locally activate the tumor microenvironment, SLR14 was delivered intratumorally (IT.) in combination with jetPEI, a cationic polymer used for in vivo transfection of nucleic acids in previous in vivo anti-tumor assays. Early administration of SLR14 at a time point when tumors are formed (day 11 post-inoculation) led to complete tumor rejection across multiple doses (1, 2, or 4 μg; Fig. 2b–d). After late administration (day 20), SLR14 significantly extended survival and induced tumor rejection in two mice (Extended Data Fig. 3a–d). Control assays with intracerebral administration of scrambled RNA, SLR14 monophosphate, or SLR14-OH induced no survival benefit, while treatments with CpG oligodeoxynucleotides resulted in a limited survival compared to SLR14 treatment (Extended Data Fig. 3e). We next administered SLR14 into the syngeneic SB28 GBM model, generated by intracerebral inoculation of non-immunotherapy responsive tumor cells ^34^. In this model, the agonist SLR14 prolonged survival of SB28 GBM-bearing mice compared to control when administered alone (Fig. 2e–h), and further enhanced survival when combined with temozolomide (TMZ), resulting in improved outcomes in comparison to either treatment alone (Extended Data Fig. 3f, g). The triple therapy with SLR14, TMZ, plus stereotactic radiotherapy (RT, Extended Data Fig. 3h) was the most efficient combinantion, resulting in long-term survival and tumor rejection in almost all treated mice (Fig. 2i-l). Therefore, SLR14 synergizes with standard therapies in syngeneic GBM-bearing mice.

**Fig. 2.**
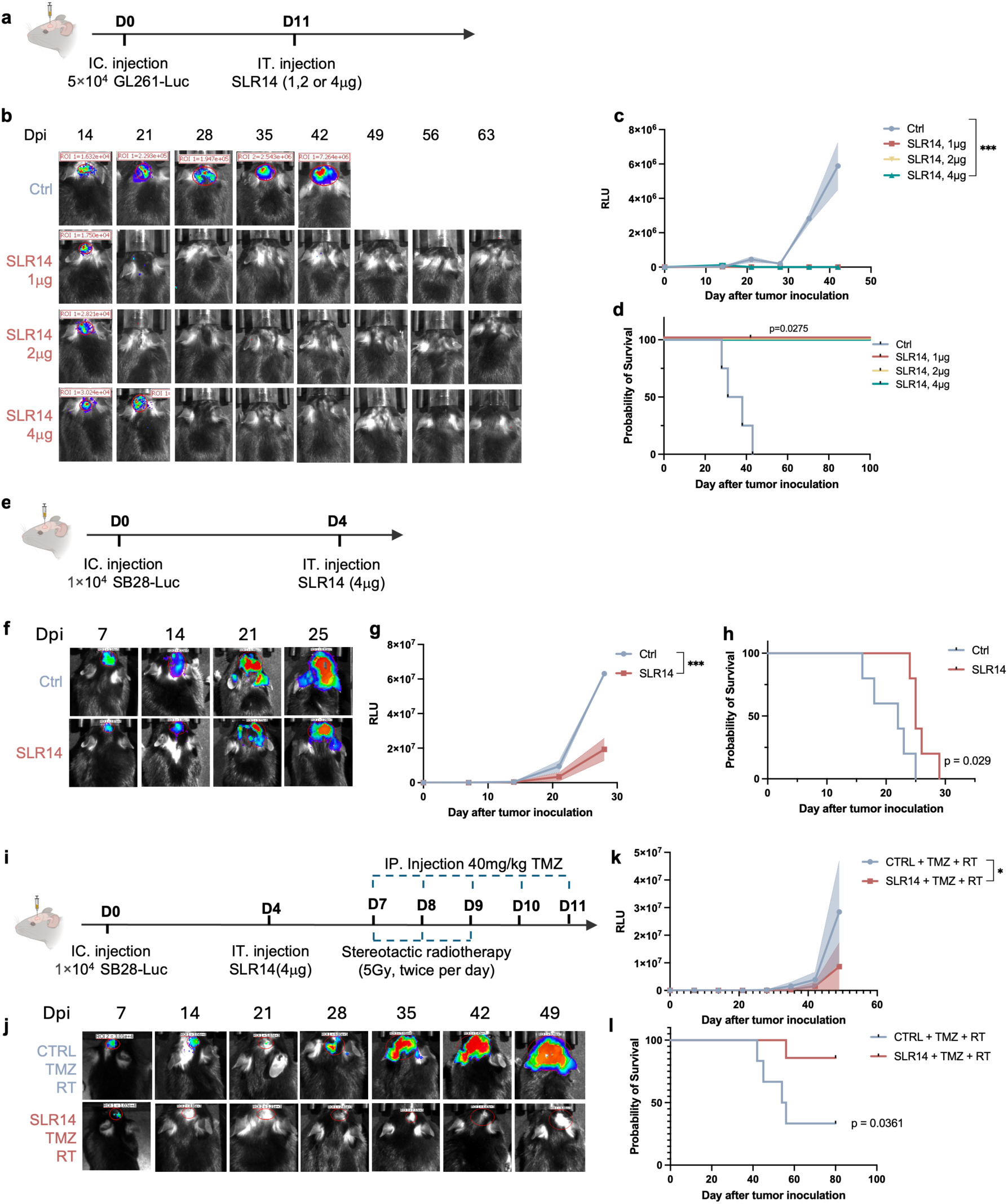
RIG-I agonist SLR14 protects against GBM and improves the outcome of standard therapies. **a**, Schematic for the experimental design of (**b-d**), using the GL261-Luc model. Mice were monitored for tumor volume and survival. **b**, Representative bioluminescence images showing GL261-Luc tumor burden. **c**, Quantification of tumor volume in GL261-Luc-bearing mice (n = 4 per group). **d**, Kaplan–Meier survival curves of GL261-Luc-bearing mice following SLR14 treatment (n = 4 per group). **e**, Schematic of mice procedure for (**f-h**), using the SB28-Luc model. **f**, Representative bioluminescence images showing SB28-Luc tumor burden. **g**, Quantification of tumor volume (n = 4 per group). **h**, Kaplan–Meier survival curves (n = 5 per group). **i**, Schematic of the treatment plan for (**j-l**), combining SLR14, TMZ, and radiotherapy (RT) in the SB28-Luc model. **j,** Representative bioluminescence images showing SB28-Luc tumor burden. **k**, Quantified tumor volumes in SB28-bearing mice receiving combined treatments (CTRL + TMZ + RT, n = 6; SLR14 + TMZ + RT, n = 7). **l**, Kaplan–Meier survival curves of SB28-Luc-bearing mice receiving combined treatments (CTRL + TMZ + RT, n = 6; SLR14 + TMZ + RT, n = 7).

## SLR14 promotes reprogramming of TAMs in GBMs

To characterize the cellular targets of SLR14 in vivo, we performed flow cytometry analyses of GL261 GBMs previously injected with a conjugated SLR14^647^ (Fig. 3a and Extended Data Fig. 4a). The majority of SLR14^647+^ cells detected at 1- and 2-days post injection (1- and 2-dpi) were CD45^+^ tumor-infiltrating lymphocytes (TILs) (∼60%) and this population was reduced to ∼30% of TILs at 7-dpi. Only a minority of CD45⁻ cells, including tumor and stromal cells, were labeled by SLR14^647^ at all stages (Fig. 3b, c and Extended Data Fig. 4b). As expected, the population of SLR14^647+^ CD45^+^ cells contained a majority (∼70%) of CD45^high^ CD11b^+^ macrophages, with smaller proportions of CD45^low^ CD11b^+^ microglia and very few T and B cells (Fig. 3d, e and Extended Data Fig. 4c). Immunolabeling analyses of GBM-bearing brains further confirmed the colocalization of SLR14^647^ with RIG-I in CD11b⁺ macrophages (Fig. 3f Extended Data Fig. 4d, e), with limited uptake by microglial cells.

**Fig. 3.**
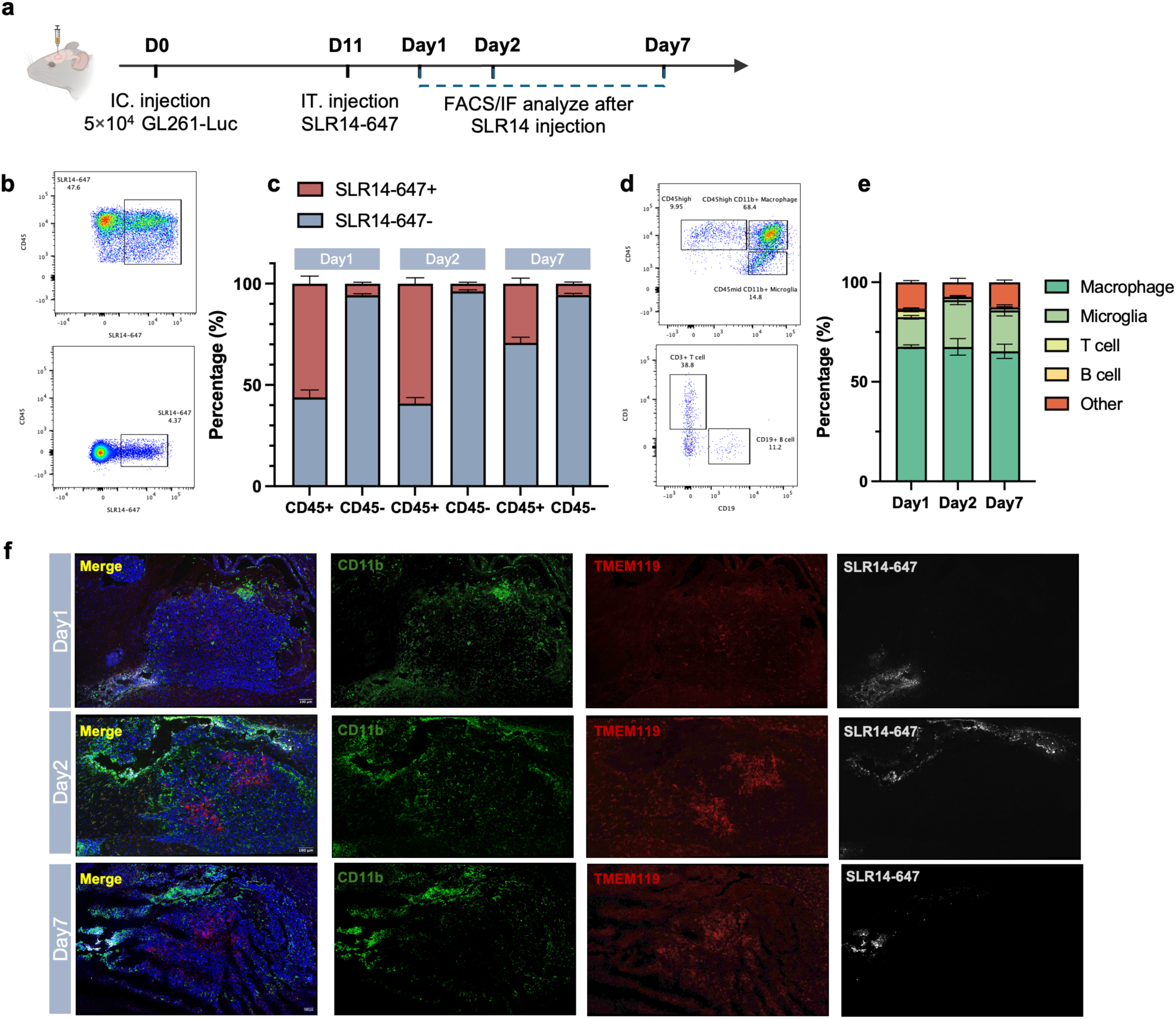
**Immunophenotype of RIG-I agonist targets after IT. injection of SLR14**^647^. **a**, Schematic of experimental design in (**b-f**). **b**, Contour plots of SLR14^647+^ cells in total CD45^+^ or CD45^-^ cells within the tumor at 1-dpi of SLR14^647^. **c**, percentage of SLR14^647+^ cells in total CD45^+^ or CD45^-^ cells within tumor at 1-, 2- and 7-dpi of SLR14^647^ (n = 5 per group). **d**, Contour plots of macrophage/microglia and lymphocyte in total SLR14^647+^ CD45^+^ cells within the tumor 1-dpi of SLR14^647^. **e**, Percentage of immune cell populations within the tumor at 1-, 2- and 7-dpi of SLR14^647^ (n = 5 per group). **f**, Representative images of immunolabeled sections of GBMs collected at 1-, 2- and 7-dpi of SLR14^647^. CD11b (green), TMEM119 (red), SLR14-647 (gray), and DAPI (blue). Scale bar, 100μm.

As the RIG-I pathway activates IFN-I signaling, SLR14 is expected to increase the expression of IFN-I signaling pathway genes in TAMs. This prediction was confirmed by scRNA-seq analysis of GL261 tumors isolated at 7-dpi with SLR14 or vehicle (Fig. 4a). Gene ontology analysis of macrophages revealed significant enrichment of pathways related to the innate immune response, type I and type II IFN signaling after SLR14 treatment (Fig. 4b, c and Extended Data Fig. 5a, b). A broad set of IFN-stimulated genes (*Isg15*, *Ifit1*, *Irf1*, *Cxcl10*), antigen-presentation molecules (MHC class I genes including *H2-K1*, *H2-D1*, *H2-Q7*, *H2-Q6*, *H2-Q4*), and regulators of monocyte recruitment and macrophage function (*LY6c2*, *Ccl2*, *Ccl7*, *Il15*, *Tnf*, *S100A6*) were significantly upregulated. Noteworthy, genes associated with pro-tumor macrophage functions (*Mrc1*, *Apoe*, *Il1b*, *Il1a*) were downregulated (Fig. 4d). Five distinct populations of macrophages showed transcriptional profiles consistent with our previous analyses (Fig. 4e, f and Extended Data Fig. 5c). RIG-I-related genes (*Cxcl10*, *Isg15*, *Ifit1*, *Ifit3*) were markedly upregulated within *Rigi*⁺ TAMs after SLR14 treatment, with a stable representation of this cluster compared to control conditions (Fig. 4g-i and Extended Data Fig. 5d). Notably, expression of *Mrc1* and *Il1b*, previously implicated in immune suppression, was decreased (Extended Data Fig. 5d). Flow cytometry analysis corroborated these findings, revealing a significant increase in CD86⁺CD206⁻ M1 macrophages, and a decrease in CD86⁻CD206⁺ M2 macrophages after SLR14 treatment. Consequently, the M1/M2 ratio was markedly elevated compared with vehicle controls (Fig. 4j, k and Extended Data Fig. 5e). Immunostaining further revealed a reduction in CD206⁺ macrophage infiltration within SLR14⁺ regions (Extended Data Fig. 5f). Consistent with these results, ELISA of tumor lysates further demonstrated increased levels of the RIG-I-related chemokine CXCL10 and antitumor cytokines including IFN-γ, MCP-1, and TNF-α, alongside reduced levels of proinflammatory cytokines such as IL-6 and M-CSF (Fig. 4l). Serum VEGF level was also elevated after SLR14 treatment, but without causing leakage of the blood brain barrier (Extended Data Fig. 5g). This set of studies demonstrates SLR14 potential to repolarize TAMs toward a pro-inflammatory and antitumor phenotype.

**Fig. 4.**
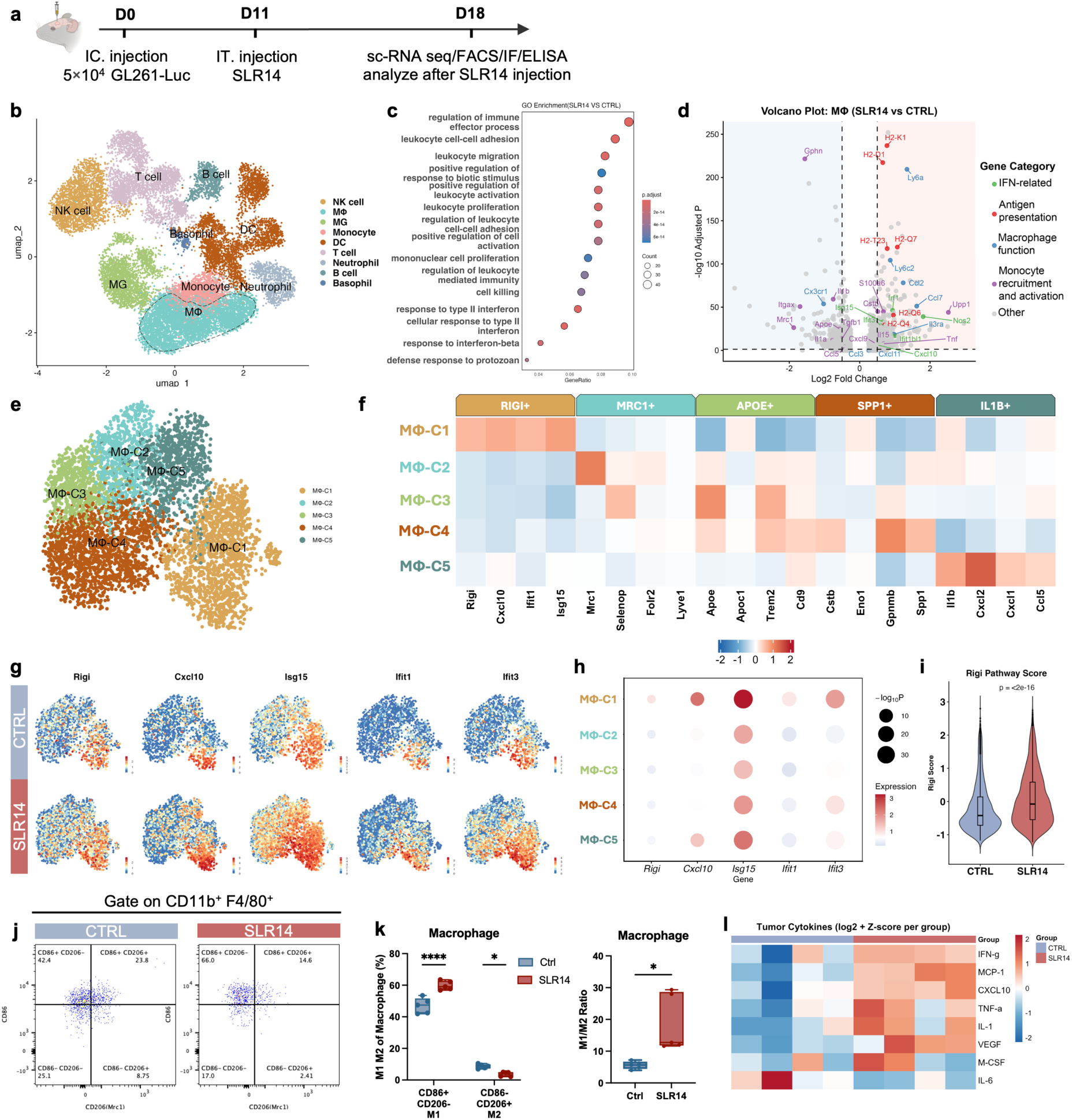
ScRNA-seq analysis of tumor immune microenvironment following SLR14 treatment. **a**, Schematic of experimental design for (**b-l**). **b,** UMAP of scRNA-seq data from GL261-bearing mouse (n = 6). **c**, Gene Ontology (GO) enrichment analysis showing the top 15 upregulated pathways in TAMs from SLR14-treated mice compared to control (n = 3 per group). **d**, Volcan plot showing significantly upregulated and downregulated genes in TAMs after SLR14 treatment (n = 3 per group). **e**, UMAP visualization of macrophage subclusters. **f**, Expression (scRNA-seq) of selected genes belonging to the indicated categories in TAM subsets. **g**, UMAP of RIG-I pathway related genes expression (*Cxcl10, Rsad2, Isg15, Ifit1, Ifit2, Ifit3)* in Mφ-C1, comparing control and SLR14-treated groups. **h**, Dot plots showing expression levels and significance (–log₁₀ P values) of RIG-I pathway-related genes in SLR14-treated mice, compared with controls. **i**, Mean expression of the *Rigi*-expressing TAM gene signature in SLR14-treated mice, compared to control. **j, k,** Contour plots (**j**) and frequency (**k**) of macrophage polarization according to CD86 and CD206 expression in tumors from control and SLR14-treated mice (n = 5 per group). **l.** Heatmap of cytokine levels measured by multiplexed bead-based ELISA from GL261 allografts at 7-dpi of SLR14 (n = 4 per group).

## Macrophage reprogramming enhances T cell infiltration and antigen presentation

We next investigated how RIG-I–mediated macrophage reprogramming influences myeloid cell interactions with the adaptive immune system within the GBM microenvironment. We first analyzed the lymphoid compartment of syngeneic GBM models. Unsupervised graph-based clustering revealed four major cell clusters manually annotated by the expression of marker genes (Fig. 5a, b). Interactions between the myeloid and lymphoid cell clusters were predicted using CellChat^35^. Macrophages were found to provide the strongest chemokine output signaling to CD8^+^ T cells and NK cells (Fig. 5c). Interestingly, myeloid–lymphoid interactions involving *Cxcl9, Cxcl10*, and *Cxcl11* signaling through the *Cxcr3* receptor on T, NK, and NKT cells were predominantly predicted to originate from *Rigi*⁺ macrophages, relative to other myeloid cell clusters. The predicted intraction confirmed the increased probability of these chemokine signaling communications between RIG-I^+^ macrophages and T/NK/NK T cells following SLR14 treatment (Extended Data Fig. 6a). Accordingly, flow cytometry analysis confirmed increased infiltration of CD8^+^T cells and NK cells in GBMs after SLR14 treatment, although not in the deep cervical lymph nodes (dCLNs) (Fig. 5d-f and Extended Data Fig. 6b). In NK cells, SLR14 increased the expression of activation marker NCR1, indicating that cytotoxic activity was stimulated (Extended Data Fig. 6c, d). In contrast, the number of CD4⁺ T cells and B cells in the brain and dCLNs did not significantly differ between treatment conditions (Fig. 5d-f). Immunolabeling of GBM-bearing brains showed CD3⁺ T cells and NK1.1⁺ NK cells mostly localized where SLR14 had access to the tumor microenvironment (Fig. 5g). In addition, the probability of communication between MHCI molecules and CD8 T cell receptors was also higher in the macrophages (Fig. 5h). Flow cytometry analysis allowed us to access surface MHCI expression across distinct myeloid cell populations (Extended Data Fig. 6e). Macrophage and DC exhibited higher MHCI expression compared to other myeloid cells, with MHCI expression being selectively increased in macrophages following SLR14 treatment (Fig. 5i-k). MHCI–related gene expressions were also upregulated in macrophages following SLR14 treatment (Fig. 5l and Extended Data Fig. 6f, g).

**Fig. 5.**
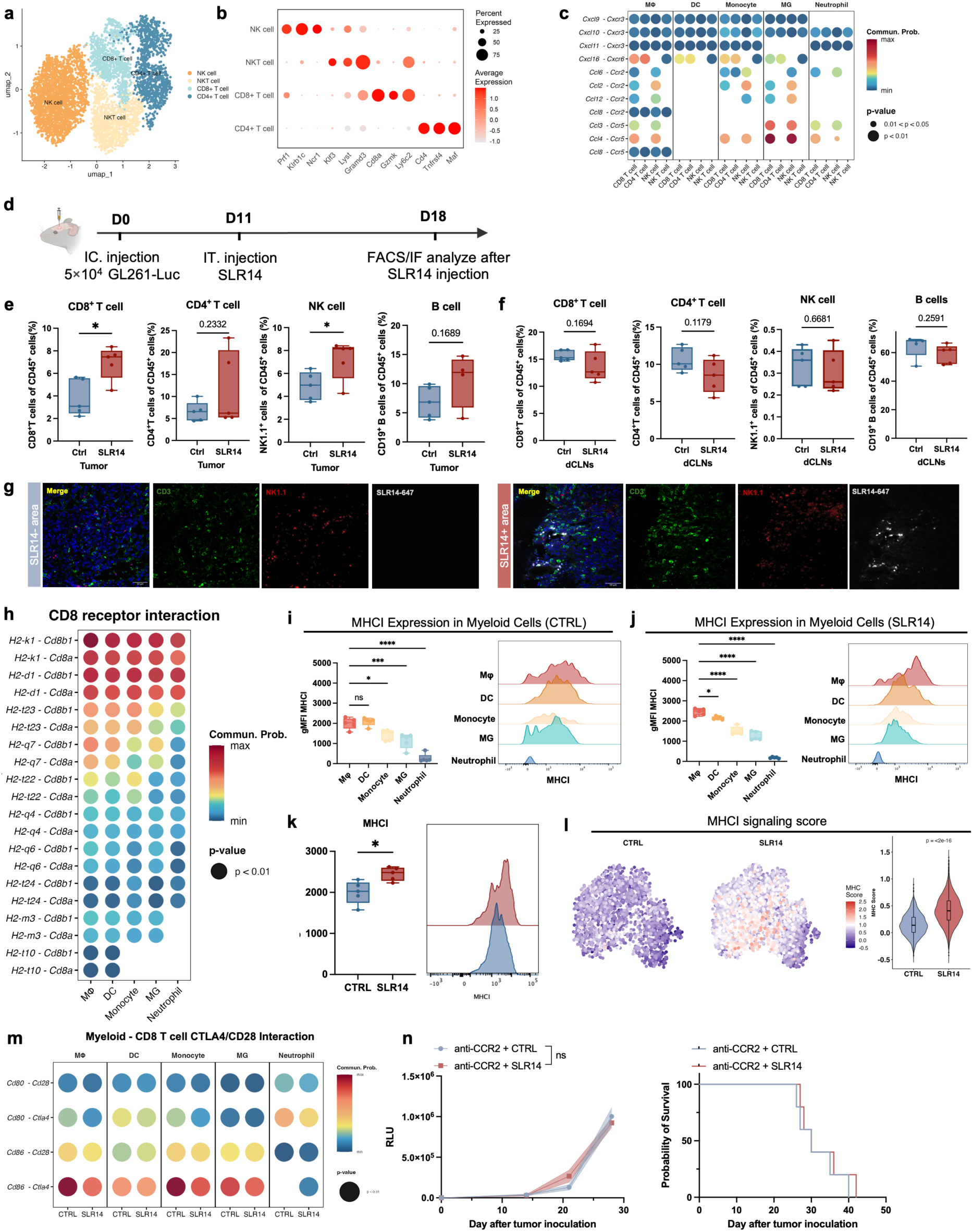
SLR14 enhances antigen presentation and cytotoxic T cell infiltration in GBM. **a**, UMAP visualization of lymphoid cell subclusters. **b**, Dot plot representing the scaled average expression of marker genes in clusters from (**a)**. **c**, Ligand-receptor pairs contributing to chemokine signaling from myeloid to lymphoid population. The dot color represents communication probability, and size indicates *p* value for interaction significance. **d**, Schematic of experimental design for (**e-m**). **e**, Quantification of the proportion of CD8^+^ T cells, CD4^+^ T cells, NK cell, and B cells in total CD45⁺ cells within the tumor (n = 5 per group). **f**, Quantification of CD8^+^, CD4^+^, NK cell, and B cell subpopulations in total CD45⁺ cells within dCLNs (n = 5 per group). **g**, Representative images of immunolabeled sections of the brain, showing NK1.1^+^ NK cells (red) and CD3^+^ T cells (green) in SLR14-negative or positive areas of the tumor. SLR14-647 (gray), and DAPI (blue). Scale bar, 50μm. **h**, Dot plot of all MHCI-related ligand-receptor pairs contributing to signaling from myeloid populations to CD8 T cells. The dot color represents communication probability, and size indicates *p* value for interaction significance. **i, j**, gMFI of MHCI of myeloid populations and representative histogram from CTRL (**i**) and SLR14 treatment (**j**) group (n = 5 per group). **k**, gMFI of MHCI of macrophage and representative histogram from control and SLR14-treated mice (n = 5 per group). **l**, UMAP embedding of single-cell profiles colored by MHCI score in macrophages (left) and a violin plot showing the distribution of MHCI score per macrophage cells in each condition (right). **m,** Dot plot of CD86/CD80– CD28/CTLA4 related ligand-receptor pairs contributing to signaling from myeloid populations to CD8 T cells. The dot color represents communication probability, and size indicates *p* value for interaction significance. **n,** Quantified tumor volumes (left) and Kaplan–Meier survival curves (right) following SLR14 treatment and anti-CCR2 depletion (n = 5 per group).

SLR14 was found to increase the macrophage expression of CD86, a costimulatory signal of T cell activation engaging either CD28 or CTLA4 to boost or dampen immune response, respectively (Extended Data Fig. 6h). Communication probability between CD86 on macrophage or monocyte and CTLA4 on CD8⁺ T cells was reduced by SLR14 treatment (Fig. 5m), suggesting reduced inhibitory signaling to T cells upon SLR14. The requirement of macrophages for the response to SLR14 was confirmed by treatment of GBM-bearing mice with an anti-CCR2 antibody that blocks monocyte trafficking to the tumor site, thereby generating a tumor microenvironment relatively deficient in monocyte-derived macrophages. The therapeutic benefit of SLR14 against GL261 GBM was markedly attenuated by anti-CCR2 antibody as compared to non-treated controls (Fig. 5n and Extended Data Fig. 6i), and SLR14-treated mice (Fig. 2d), further supporting a critical requirement for monocyte-derived macrophages in mediating the antitumor activity of SLR14.

## Adaptive immunity is required to mediate the anti-GBM effect of SLR14

We next asked whether the innate immune activation was sufficient to mediate the anti-GBM effect of SLR14. First, dCLNs of GL261GBM-bearing mice were ligated (Extended Data Fig. 7a) which interrupted lymphatic drainage from the brain and eradicated the development of an adaptive immune response against brain antigens^36–38^. dCLN-ligated mice failed to reject the GBM in the presence of SLR14 (Extended Data Fig. 7b), indicating that initiation of adaptive immunity within the draining lymph node is necessary for the anti-GBM response. Moreover, SLR14-treated tumor-bearing mice developed a long-term systemic immune memory, as re-challenging them with GBM GL261-Luc in the flank resulted in no detectable tumors (Extended Data Fig. 7c, d).

To dissect the lymphoid cell subsets involved in SLR14-induced adaptive immunity, we next carried out antibody-mediated depletion of CD4⁺ and/or CD8⁺ T cells and challenged them with GL261 GBM in the presence of SLR14 (Extended Data Fig. 7e). Depletion of CD8⁺ T cells almost completely abolished the tumor-suppressive effect of SLR14, whereas CD4⁺ T-cell depletion caused a partial loss of SLR14 effect, and combined CD4⁺ and CD8⁺ T depletion diminished the therapeutic benefit of SLR14 (Extended Data Fig. 7f, g). In contrast, additional depletion of B cells with anti-CD19 antibodies did not affect tumor growth or survival (Extended Data Fig. 7h, i). These results indicate that the antitumor activity of SLR14 mainly depends on T-cell–mediated immune responses. Consistent with our prior report in melanoma^25^, SLR14 treatment slightly extends the survival of *RAG1*^-/-^ mice, which are T and B-cell-deficient, after GBM inoculation, suggesting that another immune cell type as NK cells can contribute to the anti-tumor response to SLR14 (Extended Data Fig. 7j-l). We then depleted NK cells using anti-NK1.1 antibodies which reduced the benefit of SLR14 treatment (Extended Data Fig. 7m, n).

## SLR14 alleviates T cell exhaustion and enhances antitumor effector function

Finally, we investigated the specific T cell subsets responding to SLR14 treatment. Five CD8^+^ T cell clusters (Fig. 6a) were annotated based on their top differentially expressed genes and the expression of established exhaustion-associated markers, including *Tcf7*, *Slamf6*, *Havcr2*, and *Pdcd1*. Naïve-like T cells (Tnaive_like) and progenitor exhausted T cells (Tex_prog) were characterized by the expression of memory and naïve markers such as *Tcf7*, *Sell*, and *Ccr7*, with Tex_prog additionally exhibiting higher expression of *Slamf6* and *Pdcd1*. Effector T cells (Teff) displayed an activated phenotype marked by elevated expression of *Nr4a1*, *Cd44*, and *Cx3cr1*. Intermediate exhausted T cells (Tex_int) expressed high levels of *Ccl5*, *Gzmk*, and *Cxcr3*, accompanied by increased *Pdcd1* expression. In contrast, terminally exhausted T cells (Tex_term) exhibited the highest T cell exhaustion signature score (Fig. 6b) and the strongest expression of exhaustion-associated genes, including *Pdcd1*, *Lag3*, and *Havcr2*, together with a loss of *Tcf7* expression (Extended Data Fig. 8a-c). Moreover, CD8⁺ T cells exhibited reduced expression of exhaustion-associated gene signatures following SLR14 treatment (Fig. 6c). Consistently, CD8⁺ T cells from SLR14-treated tumors were enriched for gene signatures associated with Tex_prog (Fig. 6d), whereas Tex_term–associated signatures predominated in control tumors (Fig. 6e).

**Fig. 6.**
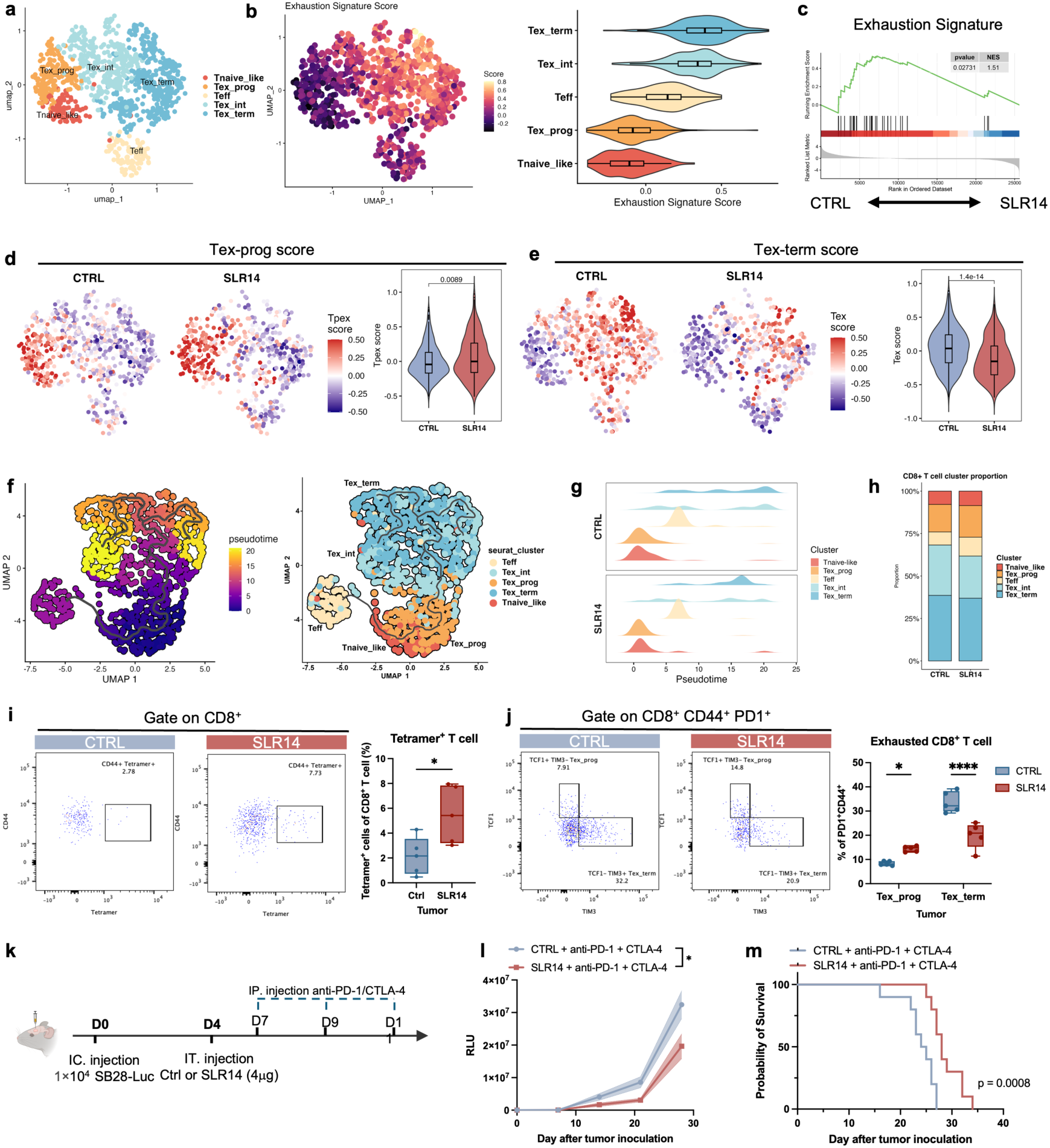
SLR14 alleviates T cell exhaustion and synergizes with immune checkpoint therapy in GBM. **a**, UMAP visualization of CD8 T cell subclusters. **b**, UMAP embedding of single-cell profiles colored by exhaustion score in CD8 T cell (left) and a violin plot showing the distribution of exhaustion score in each cluster(right). **c**, Gene set enrichment analysis (GSEA) of the T cell exhaustion signature comparing CTRL and SLR14 conditions, revealing significant enrichment of the exhaustion program in CTRL compared with SLR14 (NES = 1.51, nominal p = 0.027). **d**, UMAP embedding of single-cell profiles colored by Tex-prog score in CD8 T cell (left) and a violin plot showing the distribution of Tex-prog score in each condition(right). **e,** UMAP embedding of single-cell profiles colored by Tex-term score in CD8 T cell (left) and a violin plot showing the distribution of Tex-term score in each condition(right). **f**, Trajectory analysis of CD8 T cell colored by pseudotime (left) and annotated by subclusters (right). **g,** Density distributions of CD8^+^ T cell states along pseudotime in CTRL (top) and SLR14 (bottom) conditions. **h**, Frequency of each cluster in CTRL and SLR14 treatment group. **i**, Contour plots (left) and frequency (right) of tetramer^+^ T cells in CD8^+^ T cells within the tumor (n = 5 per group). **j**, Contour plots (left), and frequency (right) of TCF1^+^TIM3^+^ Tex-prog and TCF1^+^TIM3^+^ Tex-term cells in CD8^+^ CD44^+^ PD1^+^ T cells within tumor (n = 5 per group). **k**, Schematic of the treatment plan for (**l, m**), combining SLR14 and immune checkpoint blockage (ICB) in the SB28-Luc model. **l,** Quantified tumor volumes (n = 10 per group). **m**, Kaplan–Meier survival curves of SB28-bearing mice receiving combined treatments (n = 10 per group).

We next examined lineage relationships between the T cell exhaustion clusters, using pseudotime analysis in our dataset. Two major differentiation trajectories were identified: one tracing the progression from Tnaive_like and Tex_prog through Tex_int toward Tex_term, and another leading toward Teff fate (Fig. 6f). SLR14 treatment shifted CD8⁺ T cells toward earlier pseudotime states (Fig. 6g). Importantly, the proportions of Tex_prog and Teff were increased following SLR14 treatment, whereas Tex_int and Tex_term T cell populations were reduced (Fig. 6h). Therefore, SLR14-induced immune remodeling promotes a bias of CD8⁺ T cells toward progenitor exhausted states and limits progression toward terminal exhaustion along pseudotime.

Flow cytometry analyses of tumor and dCLN cells validated that SLR14 promotes T cell priming (Extended Data Fig. 8d). Labeling with tetramers against emv2-env, an endogenous retroviral antigen overexpressed in GL261 cells^36,39^, showed an enrichment of tumor-specific CD8⁺ T cells in both the tumors and dCLNs of SLR14-treated mice (Fig. 6i and Extended Data Fig. 8e). The dCLNs exhibited a higher frequency of Tex_prog (PD-1⁺ TCF1⁺ TIM3⁻), whereas tumor sites were enriched for Tex_term (PD-1⁺ TCF1^-^ TIM3^+^) (Fig. 6j and Extended Data Fig. 8f). Notably, SLR14 specifically reduced the accumulation of terminally exhausted CD8⁺ T cells and increased the progenitor CD8⁺ T cells within tumors, without significantly altering the Tex_prog compartment in dCLNs (Fig. 6j and Extended Data Fig. 8f). In contrast, the frequency and exhaustion status of CD4⁺ T cells remained largely unchanged in both tumors and dCLNs (Extended Data Fig. 8g, h). Together with the macrophage reprogramming observed upon SLR14 treatment, these results demonstrate that SLR14 alleviates T cell exhaustion primarily through remodeling of the tumor immune microenvironment rather than by altering lymph node priming.

## SLR14 enhances immune checkpoint blockade

We interrogated a clinical dataset (GSE235676)^40^ from a trial of GBM patients treated with the anti–PD-1 antibody pembrolizumab to determine whether the response to immunotherapy may correlate with *RIGI* expression level in the tumor. The cohort included five non-responders (NRS) and seven responders (RS). Using scRNA-seq analysis, we found that *RIGI*-expressing TAMs were enriched in RS patients compared with the NRS group (Extended Data Fig. 8i, j). We inferred that elevated *RIGI* expression may promote the effects of PD1 immune checkpoint blockade and tested how much SLR14 treatment with anti-PD-1/CTL-4 antibodies improved tumor rejection and survival of GBM-bearing mice. SLR14 and anti–PD-1/CTLA-4 significantly extended survival compared with either SLR14 or checkpoint blockade alone (Fig. 6k–m). Together, these findings demonstrate that SLR14 agonism enhances immune checkpoint blockade to potentiate antitumor immunity in GBM.

## Discussion

TAMs are considered as the predominant immune population in GBM and thought to promote tumor growth, immune suppression, and resistance to therapy^41^. Our findings establish a strong correlation between the activation of a subset of *RIGI*^+^ TAMs with the priming of innate and adaptive immune cells against GBM. *RIGI*^+^ TAMs represent a subset of monocyte-derived macrophages expressing interferon-related genes in human GBMs and in murine models. *RIGI*^+^ TAMs occupy spatially restricted niches within angiogenesis–immune hubs and are associated with favorable patient outcomes, suggesting a role in maintaining local immune activity within the tumor.

In syngeneic mouse models of GBM, RIG-I activation by the agonist SLR14 induced remodeling of the tumor microenvironment to restore effective antitumor immunity and improved treatments with either chemotherapy plus radiotherapy, or anti-PD-1 immunotherapy. Intratumoral delivery of SLR14 was mainly taken up by CD11b⁺ macrophages, which induced robust type I interferon and antigen-presentation programs, while repressing immunosuppressive genes such as *Mrc1* and *Il1b*. Recent studies identified *Cxcl9*/*Cxcl10*⁺ TAMs as immunostimulatory macrophages that recruit and position T cells in immune hubs of peripheral tumors^29^, although their function in GBM remains unclear. Our data extend these observations by showing that SLR14 drives a proinflammatory and antitumor macrophage phenotype in *RIGI*⁺ TAMs, which strongly express IFN-γ, TNF-α, and CXCL10 while reducing IL-6 and M-CSF. The requirement of monocyte/macrophages for the antitumor effects of SLR14 was further supported by CCR2 blockade experiments, which abrogated the therapeutic benefit of SLR14. These findings position macrophages as essential mediators of RIG-I–driven immune remodeling in GBM and suggest that RIG-I innate immune sensing within the myeloid compartment is sufficient to reprogram an otherwise suppressive tumor microenvironment.

Several groups have highlighted the role of infiltrating myeloid cells, including TAMs^42^, DCs^43^ and microglia^44^ in shaping T cell differentiation within the tumor microenvironment. Recent work has shown that antigen presentation by TAMs in GBM drives T cells from a progenitor exhaustion state to terminal exhaustion^42^. Notably, SLR14 treatment promoted the recruitment and activation of tumor-specific T-cell populations and remodeled the intratumoral T cell compartment, as shown by the expansion of Tex prog CD8⁺ T cells and the reduction of Tex_term populations. This shift in CD8^+^ T cell phenotype was accompanied by increased expression of MHC-I and co-stimulatory molecules such as CD86 on macrophages, concomitant with enhanced antigen presentation and productive T cell engagement within the tumor.

Importantly, SLR14 treatment also increased the frequency of tumor-specific T cell populations in the dCLNs, which serve as critical sites of antigen drainage and T cell priming in GBM-bearing mice. dCLNs were enriched for progenitor exhausted CD8⁺ T cells relative to tumors, highlighting their role as an important site for the initiation of antitumor T cell responses. However, SLR14 treatment did not significantly alter the Tex_prog compartment within dCLNs, while selectively reducing terminally exhausted CD8⁺ T cells within tumors, indicating that SLR14 alleviates T cell dysfunction primarily through local remodeling of the tumor immune microenvironment rather than by altering lymph node priming. These findings suggest that RIG-I activation bridges innate and adaptive immunity within the glioma microenvironment. Functionally, depletion experiments confirmed that SLR14 efficacy relies on T-cell–mediated responses, particularly CD8⁺ T cells. Combination therapy with checkpoint inhibitors further extended survival in the otherwise resistant SB28 model, indicating that RIG-I stimulation can sensitize tumors to immune checkpoint blockade. Consistent with this, *RIGI*^+^ TAM abundance correlated with responsiveness to PD-1 blockade in human GBM datasets, implying translational relevance.

Together, these results establish RIG-I activation as a potent strategy to reprogram the glioma microenvironment and overcome its intrinsic immune suppression. By coupling innate immune activation with T cell restoration, the use of SLR14 in combination with checkpoint inhibitors, chemotherapy, or radiotherapy offers a potential therapeutic strategy against GBM.

## Methods

### Cell lines

GL261-Luc (obtained from Dr. Eric Song, Yale University) and SB28-Luc (obtained from Juan C. Vasquez, Yale University) mouse glioma cell lines were cultured in Dulbecco’s modified Eagle’s medium (DMEM)–F12 (10% FBS, 1% non-essential amino acids, 1% penicillin/streptomycin).

### Mice

Seven- to eight-week-old C57BL/6 mice and B6.129S7-Rag1tm1Mom/J mice were purchased from the Jackson Laboratory and subsequently bred and housed at Yale University. All procedures used in this study (sex-matched, age-matched) complied with federal guidelines and the institutional policies of the Yale School of Medicine Animal Care and Use Committee.

### Antibodies and tetramer

The antibodies were used for flow cytometry or immunofluorescence staining: anti-CD45 (I3/2.3, FITC, 567377), anti-CD8a (53-6.7, BV650, 563234), anti-CD3 (145-2C11, PE/Cy5,100309; 145-2C11, BV421, 100341), anti-CD11b (M1/70, BV510, 562950), anti-MRC1 (C068C2, AF700, 141734), anti-LY6G (1A8, BUV395, 563978), anti-F4/80 (BM8, PE, 123109), anti-CD11c (N41B, PE, 12-0114-82), anti-CD4 (RM4-5, AF700, 557956; GK1.5, PE/700, 100484), anti-MHCII (M5/114, BUV737, 748845), anti-CD86 (PO3, BUV496, 750437), anti-CD19 (6D5, PE/594, 115554), anti-NK1.1 (PK136, PE, 108707), anti-NCR1 (29A1.4, BV421, 562850), anti-TIM3 (7D3, BV605, 569245; 5D12, APC, 567164), anti-PD-1(EH12.1, PE/Cy7, 561272; EH12.2H7, APC/Cy7, 329921), anti-CD44 (G44-26, AF700, 56129), anti-H-2Kd-MHCI (SF1-1.1, PE/Cy7, 116621), anti-LY6C (HK1.4, BV711, 755195; AL-21, AF700, 561237), anti-TCF7 (C63D9, PE/Cy7, 90511S), Zombie NIR Fixable Viability (APC/Cy7, 423105), anti-CD11b (M1/70.15, MA5-17857), anti-TMEM119 (E3E10, 90840S), anti-MRC1 (MR503, MA5-16871), anti-CD3 (17A2, MAB4841), anti-RIG-I (PA5-20276). The emv2-env tetramer (Kb-restricted peptides aa 604–611 of p15E protein (KSPWFTTL)) was obtained from NIH tetramer core facility. The depletion antibodies anti-CD4 (GK1.5), anti-CD8 (YTS169.4), anti-PD-1 (RMP1-14), anti-CTLA4 (9H10), anti-NK1.1 (PK136) were purchased from BioXCell.

### SLR14 and other RNA constructs

The SLR14 (pp-SLR14), SLR14-mono (p-SLR14), SLR14-OH (5’-GG AUCG AUCG AUCG UUCG CGAU CGAU CGAU CC with the 5’-terminal diphosphate, monophosphate and hydroxyl, respectively), and scramble RNA(5’pp-GAAGCAAUCUCCACUUACUAGAAA) ^24^ were synthesized as previously described^25^ RNA was mixed at a ratio of 1 μg per 0.12 μl of in vivo-JETPEI (Polyplus Transfection) and vortexed for 30 s and incubated in room temperature for 15 min before use.

### Tissue processing and microscopy

Whole brain samples were dissected, fixed in 4% PFA overnight, dehydrated in 30% sucrose for 2 days. The sample were then mounted in OCT embedding compound and frozen at -20 to -80 °C. The 40 µm cryostat sections were fixed in methanol at room temperature for 10 min, washed 3 times in 0.05% PBS-Tween (PBS-T) and incubated with blocking buffer (0.05% PBS-T + 0.3% Triton X-100 + 5% BSA) at room temperature for 1 h. Sections were stained overnight at 4 °C with the following primary antibodies: Sections were washed three times in 0.05% PBS-T, and incubated with the following secondary antibodies: donkey anti-rat IgG Alexa Fluor Plus 488 (Invitrogen A48269, 1:500), donkey anti-goat IgG Alexa Fluor Plus 555 (Invitrogen A32816, 1:500), and donkey anti-rabbit IgG Alexa Fluor Plus 647 (Invitrogen A32795, 1:500). After three washes in 0.05% PBS-T, cell nuclei were stained with DAPI (Sigma-Aldrich Merck, MBD0015) at room temperature for 15 min and coverslips were mounted onto slides with. Digital images were acquired on a Lecia SP8 scanning confocal microscope using a 20× air objective or a 63× oil objective.

### Tumor inoculation

Mice were anaesthetized using a mixture of ketamine (50 mg kg^−1^) and xylazine (5 mg kg^−1^), injected intraperitoneally. Mice heads were shaved and then placed in a stereotaxic frame. After sterilization of the scalp with alcohol and betadine, a midline scalp incision was made to expose the coronal and sagittal sutures, and a burr hole was drilled 2 mm lateral to the sagittal suture and 0.5 mm posterior to the bregma. A 5-μl Hamilton syringe loaded with tumor cells was inserted into the burr hole at a depth of 3 mm from the surface of the brain and left to equilibrate for 1 min before infusion. A micro-infusion pump (World Precision Instruments) was used to infuse 1 μl of tumor cells at 0.5 μl /min. Once the infusion was finished, the syringe was left in place for another minute before removal of the syringe. Bone wax was used to fill the burr hole and the skin was stapled and cleaned. Following intramuscular administration of analgesic (meloxicam and buprenorphine, 1 mg/ kg), mice were placed in a heated cage until full recovery.

### IVIS imaging

Mice were anaesthetized using isoflurane and injected intraperitoneally with IVISbrite D-luciferin potassium (Revvity) (100 μl, 30 mg/ ml). After 10 min, mice were imaged using the IVIS Spectrum In Vivo Imaging System.

### In vivo treatments

Wild-type (WT) mice were intracranially inoculated with GL261 or SB28 cells. For SLR14 treatment, mice were anaesthetized using a mixture of ketamine (50 mg/kg) and xylazine (5 mg/kg), injected intraperitoneally. Mice heads were shaved and then placed in a stereotaxic frame same as tumor injection. A 5 μl Hamilton syringe loaded with SLR14 or control was inserted into the burr hole at a depth of 3 mm from the surface of the brain and left to equilibrate for 1 min before infusion. A micro-infusion pump (World Precision Instruments) was used to infuse 5 μl of SLR14 at 1 μl /min. Once the infusion was finished, the syringe was left in place for another minute before removal of the syringe. Bone wax was used to fill the burr hole, and the skin was stapled and cleaned. Following intramuscular administration of analgesic (meloxicam and buprenorphine, 1 mg/ kg), mice were placed in a heated cage until full recovery. GL261-bearing mice were received SLR14 treatment at day 11 post tumor inoculation, SB28-bearing mice were received SLR14 treatment at day 4 post tumor inoculation. For chemotherapy, mice received intraperitoneal injections of temozolomide (TMZ; 40 mg per mouse) beginning on day 7 post-inoculation, administered once daily for five consecutive days. For stereotactic radiotherapy, mice were anesthetized with isoflurane and scanned by micro-CT to localize tumor sites. Normal brain tissue and tumor regions were delineated to ensure targeted irradiation. A 2 mm beam was used to deliver 5 Gy per exposure to the tumor site, administered twice daily (total 10 Gy per day) for three consecutive days. For immunotherapy, mice were injected intraperitoneally with anti–PD-1 (200 μg per mouse) and anti–CTLA-4 (200 μg per mouse) antibodies starting on day 7, with injections repeated every other day for a total of three doses. For NK cell depletion, 200 µg anti-NK1.1 (clone PK136, BioXcell) was administered every 4 days through i.p. injections, starting at day 1 before tumor induction. For T cell depletion, 200 µg anti-CD8 or anti-CD4 was administered every 4 days through i.p. injections, starting at day 1 before tumor induction. For B cell depletion, 200 µg anti-CD19 (clone X, BioXcell) was administered every 4 days through i.p. injections, starting at day 1 before tumor induction. Whereas control mice were treated with isotype control (rat IgG2b). All cell depletion was confirmed by flow cytometry.

#### Ligation of deep cervical lymph nodes

For ligation of lymph nodes, mice were anaesthetized using ketamine and xylazine, and the rostral neck was shaved and disinfected. A 2-cm incision was made and the salivary glands containing the superficial cervical lymph nodes were retracted and the deep cervical lymph nodes were visualized. The afferent lymph vessels were tied off with a 4-0 Vicrylsuture and then cauterized. The incision was closed with a 4-0 Vicrylsuture and mice were subjected to the same postoperative procedures as above.

#### Flank tumor inoculation

Mice were anaesthetized using ketamine and xylazine, and the flank was shaved and disinfected. A 1-ml syringe with a 30G needle was used to deliver 100 μl of 500,000 cells subcutaneously. For GL261-Luc cells, cells were mixed in a 1:1 volume with Matrigel (Corning) before delivering.

### Isolation of immune cells and flow cytometry

Tissues (brain or lymph nodes) were collected and incubated in a collected and incubated in a digestion cocktail containing 1 mg/ml collagenase D (Roche) and 30 μg/ml DNase I (Sigma-Aldrich) in complete RPMI (10% FBS) at 37 °C for 30 min. Tissue was then filtered through a 70-μm filter. For brain tissues, cells were mixed in 4 ml of 25% Percoll (Sigma-Aldrich) solution and centrifuged at 530 g for 15 min without a brake. The Percoll layer was removed, and cells were diluted in 5 ml of 1% BSA. Single-cell suspensions were incubated with mouse FcγIII/II receptor (CD16/CD32) blocking antibody for 10 min on ice and pelleted by centrifugation. Cell viability was assessed by Zombie NIR Live/Dead staining, applied for 30 min at 4 °C. Surface or tetramer staining was then performed with fluorophore-conjugated primary antibodies for 30 min at 4 °C. Cells were washed to remove excess antibodies and resuspended in 1% BSA for multiparameter analyses on the LSR Fortessa flow cytometer (Becton Dickinson), and then analyzed using FlowJo software.

#### Bulk RNA-seq analyze from TCGA

Bulk RNA-seq expression data and clinical annotations were obtained from The Cancer Genome Atlas (TCGA) and the Genotype-Tissue Expression (GTEx) project via the UCSC Xena platform. TCGA primary tumor samples were used as tumor specimens, and GTEx normal tissue samples were used as controls.

Gene expression levels were quantified as transcripts per million (TPM) based on RSEM normalization and were log₂-transformed where indicated. For pan-cancer analyses, GTEx normal tissues were matched to TCGA tumor samples according to primary anatomical site. For glioma analyses, including Brain Lower Grade Glioma (LGG) and Glioblastoma Multiforme (GBM), normal controls were restricted to cortical brain regions, specifically GTEx Brain – Cortex. LGG and GBM were analyzed as separate cancer types using the same set of cortical normal brain controls. Expression differences between tumor and normal samples were assessed using two-sided Wilcoxon rank-sum tests. Effect sizes were calculated as differences in median expression levels. P values were adjusted for multiple testing using the Benjamini–Hochberg false discovery rate method.

#### Sample preparation for scRNA-seq

For scRNA-seq experiments to characterize the tumor microenvironment, 3v3 (CTRL or SLR14) tumors were isolated at day 18 after injection (7 day after SLR14 treatment) and were processed as described above. The CD45^+^ live fraction was isolated by FACS, and approximately 1 × 10^5^ cells were collected. For all the scRNA-seq samples, a Chromium GEM OCM Single cell 5′ kit with Dual Index was used according to the manufacturer’s instructions. GEMs were generated on Chromium X (10x Genomics) with a target of 20,000 cells recovered, and libraries prepared according to the manufacturer’s instructions (CG000527, 10x Genomics). Sequencing was performed on NovaSeq PE100 (Illumina) with a target of 20,000 reads per cell.

#### scRNA-seq analysis of GBM

Seq samples were pre-processed using Cell Ranger Multi (v.7.1.0). Cell Ranger-filtered feature–barcode matrices were further analyzed by Seurat (v.5.2.0) in RStudio (v.4.5.1) and further filtered to retain cells with more than 400 detected genes, and less than 10% of mitochondrial. Data were normalized and scaled by SCT method, and dimensionality reduction was performed using principal component analysis on the top 3,000 most variable genes. Harmony was used for the integration of cells from different samples. The first 30 Harmony dimensions were used to identify immune cell subclusters with a resolution of 0.5 that were further assigned to cell types using known markers and publicly available myeloid reference datasets. Wilcoxon rank-sum test implemented in Presto (v.1.0.0) was used to identify differentially expressed genes (DEGs). *RIGI*^+^ TAM scores, MHCI score, Exhaustion signature score, Tex_prog score, and Tex_term score were calculated using the “AddModuleScore” function of Seurat.

#### Trajectories analysis

To understand differentiation trajectories of myeloid cells within the tumor microenvironment, we performed analysis of the monocyte and macrophage compartment by monocle3^45^. The monocle3 data were combined with the scRNA-seq object that had been filtered to keep the data for monocyte and macrophage populations (monocyte, *Rigi*⁺ TAMs, *Mrc1*⁺ TAMs, *Apoe1*⁺ TAMs, *Spp1*⁺ TAMs, and *Il1b*⁺ TAMs) for each condition (CTRL or SLR14). To understand differentiation trajectories of CD8^+^ T cells, CD8^+^ T cells were reclustered and analyzed by monocle3.

#### Analysis of publicly available GBM datasets

For mouse GBM sample data from previously published datasets (Gene Expression Omnibus (GEO) accession numbers GSE193526, GSE264251, and GSE215239), the matrix data were pre-processed as described in the corresponding publications. Batch correction and integration across samples were performed using Harmony, and clustering was recalculated with a resolution parameter of 1.0. For human GBM (hGBM) datasets, single-cell RNA-seq data from eight newly diagnosed GBM patients (GEO accession GSE182109) were analyzed to characterize transcriptional heterogeneity. In addition, data from GSE235676, comprising 24 GBM patients, including those who received neoadjuvant therapy, were analyzed for comparative and clinical correlation studies.

#### Spatial transcriptomics of human GBM samples

Single-cell spatial transcriptomics and biopsy sample data were obtained from previously published Dryad database (https://datadryad.org/stash/dataset/doi:10.5061/dryad.h70rxwdmj)^30^. Data processing and visualization were performed using SPATA2 (v3.1.3) in RStudio (v4.5.1). Spatial state layers were clustered and annotated according to a previously published framework describing the multi-layered organization of glioblastoma^46^.

#### TCGA data and deconvolution analysis

The correlation of OS and gene expression from TCGA GBM datasets were obtained from GEPIA2 (http://gepia.cancer-pku.cn/). The high and low expression or proportion was grouped based on most fitted percentiles.

#### Cytokine analysis by Luminex assay

To quantify cytokine levels within tumor tissues, a multiplex Luminex assay (LXSAMSM-10) was performed to detect ten cytokines according to the manufacturer’s instructions. Briefly, 50 μl of standards or samples were added to each well, followed by 50 μl of diluted microparticle cocktail. Plates were incubated for 2 h at room temperature on a shaker (800 rpm), then washed three times with 100 μl of wash buffer. Next, 50 μl of diluted biotin–antibody cocktail was added and incubated for 1 h at room temperature with shaking, followed by washing as above. Wells were then incubated with 50 μl of diluted streptavidin–PE for 30 min, washed again, and resuspended in 100 μl of wash buffer for 2 min at room temperature. Plates were read within 90 min using a Luminex.

#### Quantification and statistical analyses

Results are illustrated as mean ± s.d. or mean ± s.e.m. Graphs show data from at least two independent repeats. Significance was defined as *: p < 0.05, **: p < 0.01, ***: p < 0.001. Statistical analysis was conducted either using GraphPad Prism v10.0 (GraphPad Software) or RStudio (v4.5.1). Statistical tests, exact value of n and what n represents are mentioned in the figure legends. Tumor growth curves were analyzed using two-way repeated-measures ANOVA with time and treatment as factors. Survival curves were generated using the Kaplan–Meier method and compared using the log-rank (Mantel–Cox) test. Statistical tests with adjustment for multiple comparisons and exact P values are reported in the Source Data.

## Code availability

This study did not generate custom code for the analyses. Standard workflows and open-source R packages and software were used (Methods).

## Acknowledgements

We thank members of the A.E. and J.L.T. laboratories for insightful discussions and assistance with experimental protocols; P.L. for help with flow cytometry panel design; O.F. for providing materials; and Phil Coish for helpful comments on the manuscript. This work was supported by the National Institutes of Health (R01NS130057) and the Yale Cancer Center–The Chenevert Family Brain Tumor Center at Yale (H.X., S.L., F.L., A.E., J.L.T.). Additional support was provided by the Howard Hughes Medical Institute (A.I., A.M.P.), Yale School of Medicine (A.I., A.M.P., and E.S.), and Fondation ARC (grant PAIR TUMC21-001), the Paris Brain Institute (ICM), and Paris Brain Institute America (PBIA) (M.T.).

## Author contributions

H.X., A.I., A.M.P and J.L.T. planned the project. H.X. and J.L.T designed, analysed and interpreted data and wrote the manuscript. H.X., S.L. and F.L. performed experiments and analysed data. J.-L.T. provided AAV material.,P.L, and O.F. provided expertise, materials and analysis of data.

## Competing interests

A.I. and A.M.P. co-founded RIGImmune. A.I. and E.S. co-founded Rho Bio. A.I. co-founded Xanadu Bio and PanV, and is a member of the Board of Directors of Roche Holding Ltd and Genentech.

---

**Extended Data Fig. 1.**
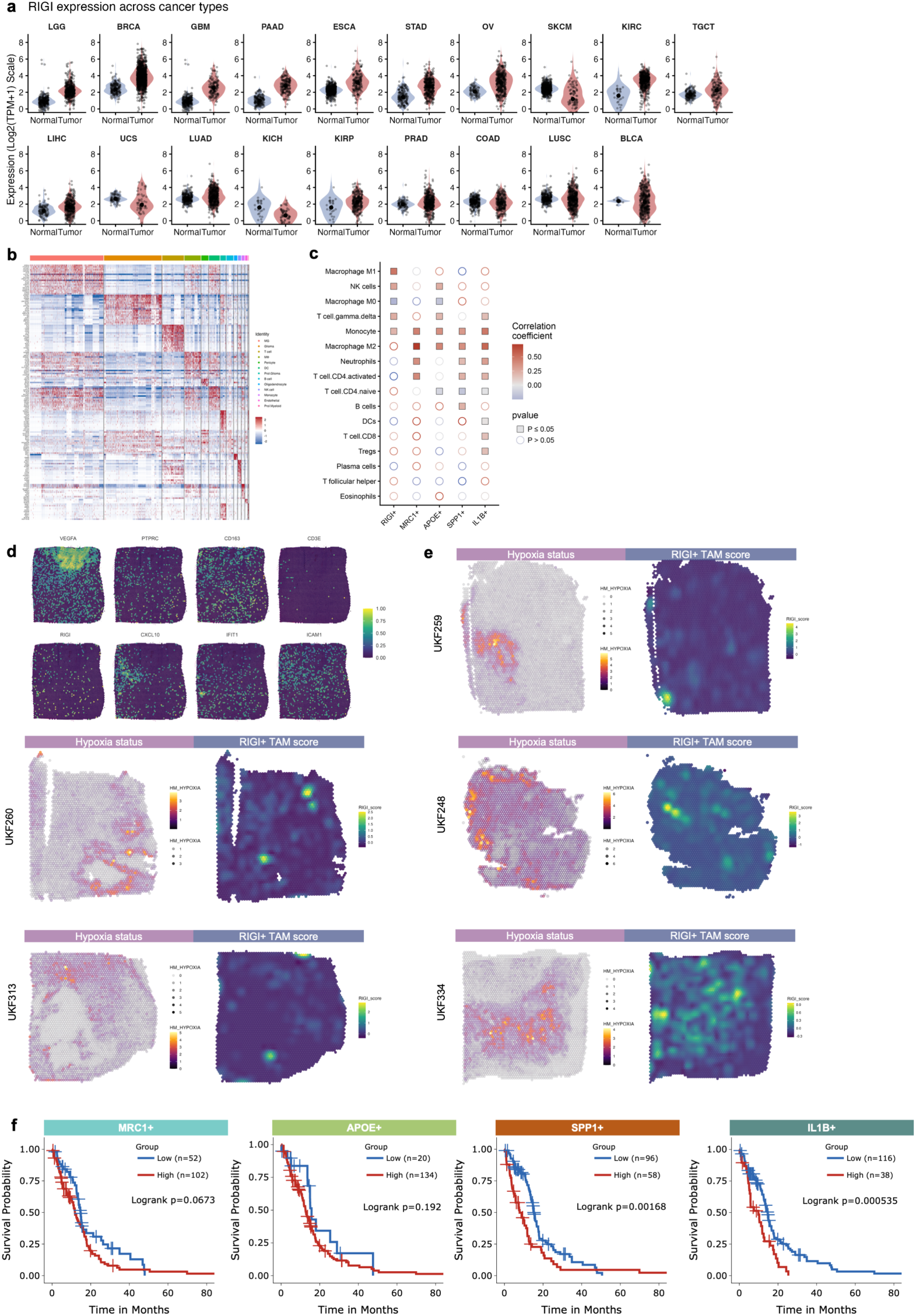
scRNA-seq and spatial transcriptomics analyses of human GBMs. **a**, Violin plot showing *RIGI* gene expression across different cancer types and their origin tissue (data from TCGA tumor and GTEx normal). **b**, Heatmap of scaled expression of the top 25 marker genes for each scRNA-Seq cluster. **c**, Correlation between five macrophage cluster marker genes expression and representative immune-cell-population marker genes in TCGA GBM patients. **d**, Marker genes of *RIGI*-expressing TAMs at spatial resolution from case UKF275 of the Dryad database. **e,** Surface plots showing the transcriptional profiling of hypoxia score (left) and *RIGI*-expressing TAMs score (right) in human GBM tissue (n = 5, public dataset from the Dryad database). **f,** Survival probability of patients with GBM (from TCGA data) stratified with respect to the *MRC1*^+^, *APOE*^+^, *SPP1*^+^, *IL1B*^+^TAM gene signature.

**Extended Data Fig. 2.**
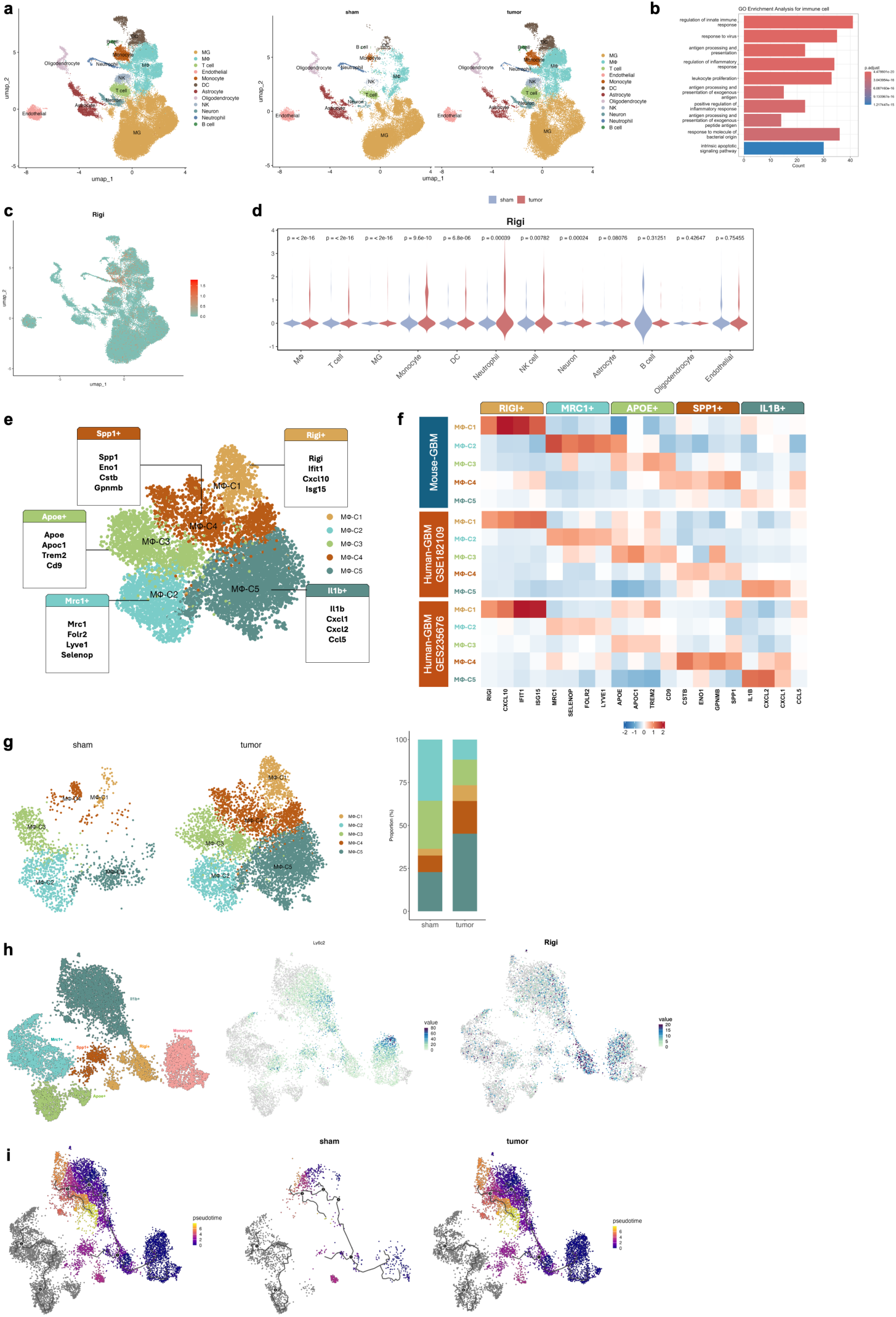
Single-cell RNA-seq analysis of *Rigi* expression in the brain of control and GBM-bearing mice. **a**, UMAP of scRNA-Seq of all cells (left) and split by sham (n = 5) and tumors (n = 6) (right) from mouse GBM (GSE193526, GSE264251 and GSE215239). **b**, GO enrichment analysis showing upregulated pathways in tumor-infiltrating immune cells, including both innate and adaptive immune responses. **c**, UMAP visualization of *Rigi* gene expression across all cell types. **d**, Violin plot showing upregulated *Rigi* expression in tumor infiltrating immune cells. **e**, UMAP of scRNA-seq data from macrophages. **f**, Expression (scRNA-seq) of selected genes belonging to the indicated categories in TAM subsets across different mouse and human database. **g**, UMAP showing clustering (left) and bar plots showing frequencies (right, scRNA-Seq) of the indicated macrophage types in either sham or tumor tissue. **h**, Sub clustering of macrophages and monocytes (left) and corresponding expression (scRNA-seq) of *Ly6c2* and *Rigi* (right). **i**, Pseudo time trajectory analysis of macrophages and monocytes (left) and comparison between sham (n = 5) and tumor-bearing (n = 6) mice (right).

**Extended Data Fig. 3.**
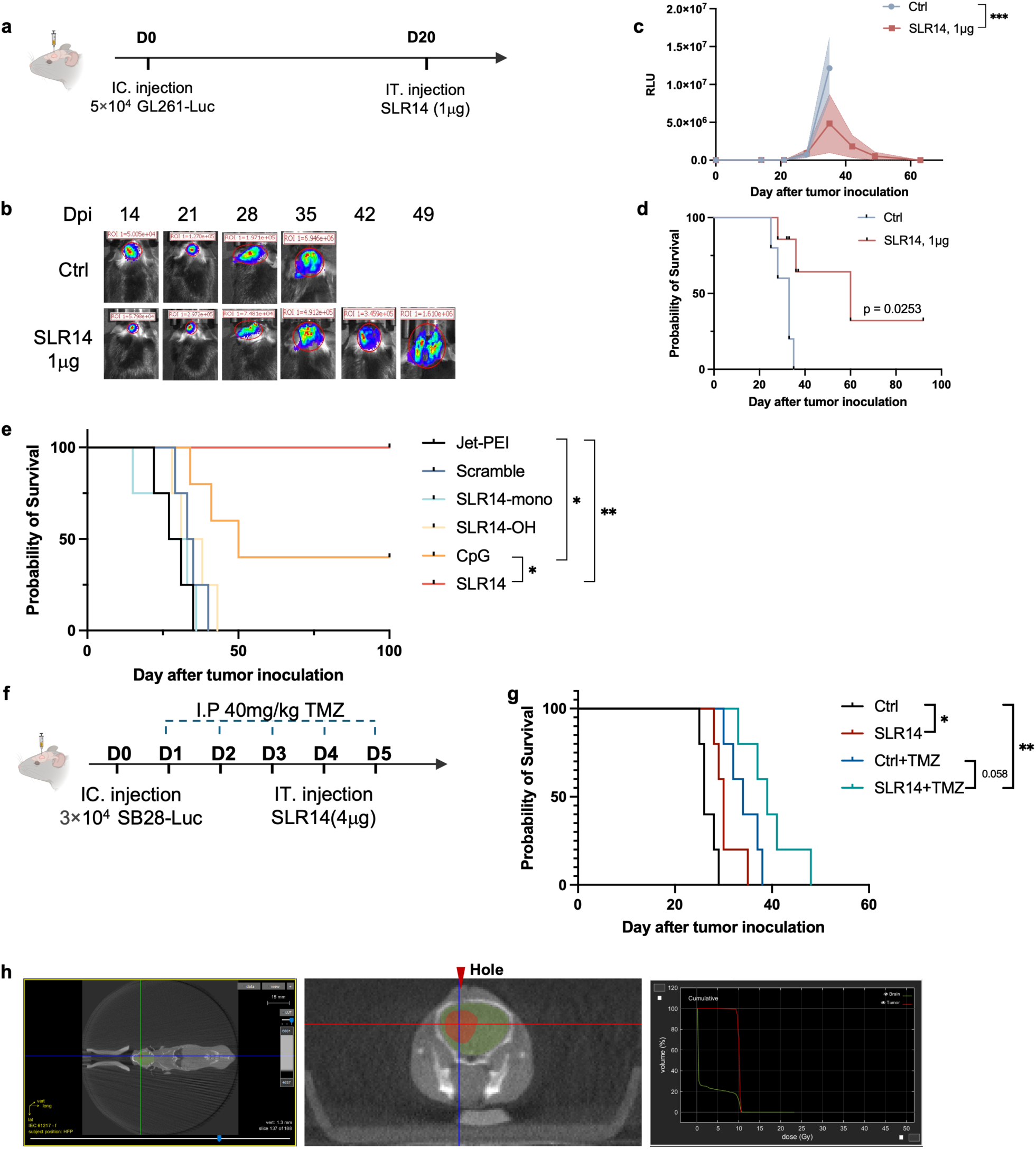
SLR14 demonstrates therapeutic effects in late-stage GL-261. **a**, Schematic for experimental design in (**b-i**), using the GL-261 model. **b**, Representative bioluminescence images showing GL261 tumor burden. **c**, Quantification of tumor volume (n = 5 per group). **d**, Kaplan–Meier survival curves (n = 5 per group). **e,** Kaplan–Meier survival curves comparing different treatments, including SLR14-monoCpG (4 μg), Scramble (4 μg), SLR14-OH (4 μg), CpG (4 μg) and SLR14 (4 μg). **f**, Schematic of the experiment in (**g**) combining treatment regimen with temozolomide (TMZ) alone or in combination with SLR14 in SB28 GBM-bearing mice. **g**, Kaplan–Meier survival curves (n = 5 per group). **h**, Scheme showing the stereotactic radiotherapy strategy. Tumor sites were identified by the syringe hole in the skull cap and received a focused 10 Gy dose, while surrounding brain tissue received minimal radiation exposure.

**Extended Data Fig. 4.**
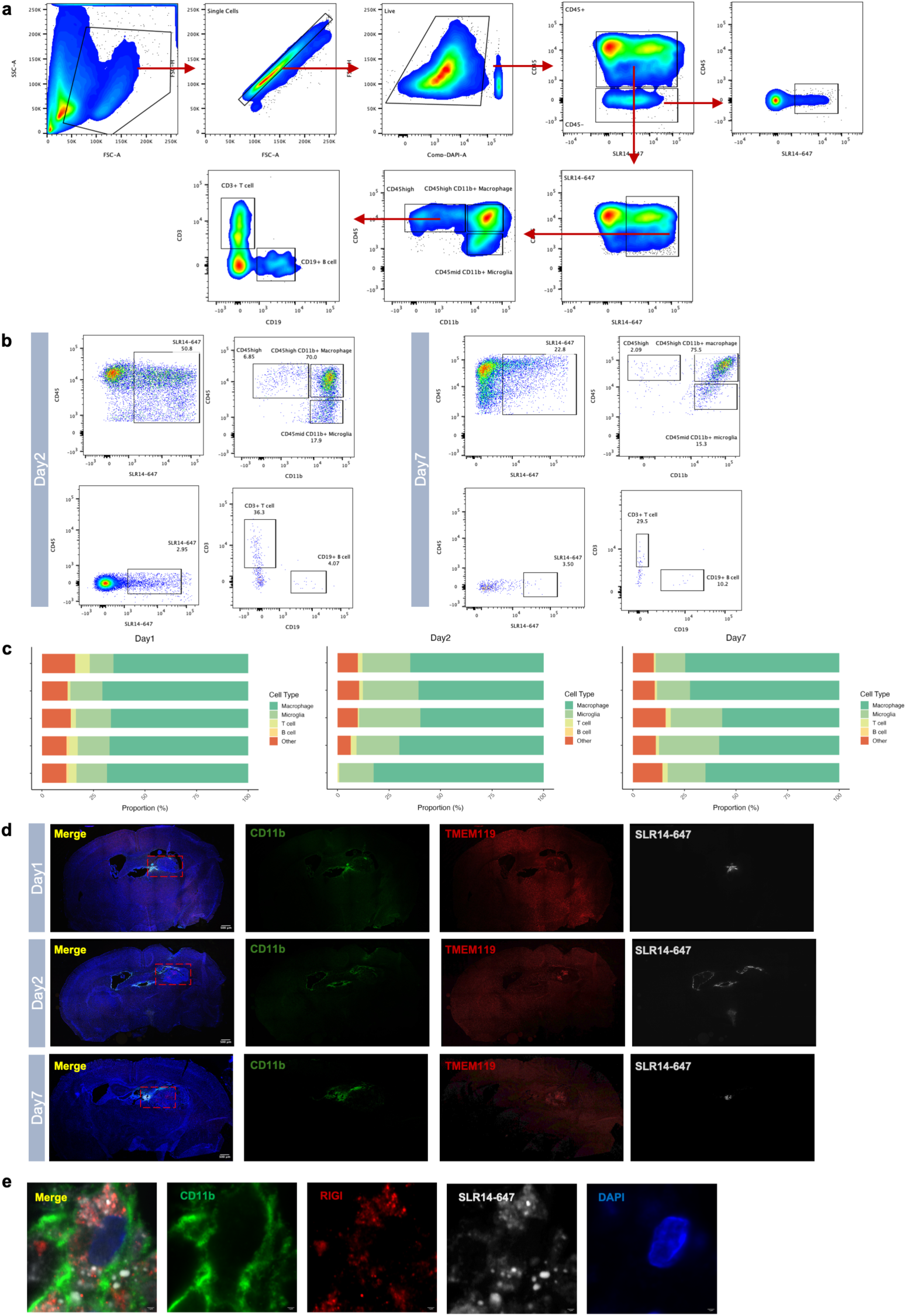
Flow cytometry strategy and immunofluorescence staining of immune cell populations after SLR14-647 treatment. **a**, Gating strategy for macrophage, microglia, and lymphocyte analyses. **b**, Contour plots of SLR14^+^ cells in total CD45^+^ or CD45^-^ cells, and macrophage/microglia and lymphocyte in total SLR14^+^ CD45^+^ cells within the tumor at 2-dpi (left) and at 7-dpi (right). **c**, Frequency (right) of macrophage, microglia and lymphocyte in CD45^+^ cells within the tumor (n = 5 per group). **d,** Representative image of whole-brain sections collected at 1-, 2-, and 7-dpi of SLR14^647^ and immunolabeled for detecting cells expressing CD11b (green), TMEM119 (red), SLR14-647 (gray), and DAPI (blue). Scale bar, 500μm. **e**, Representative image of immunolabeled tumors collected at 7-dpi of SLR14^647^. CD11b (green), RIGI (red), SLR14-647 (gray), and DAPI (blue). Scale bar, 1μm.

**Extended Data Fig. 5.**
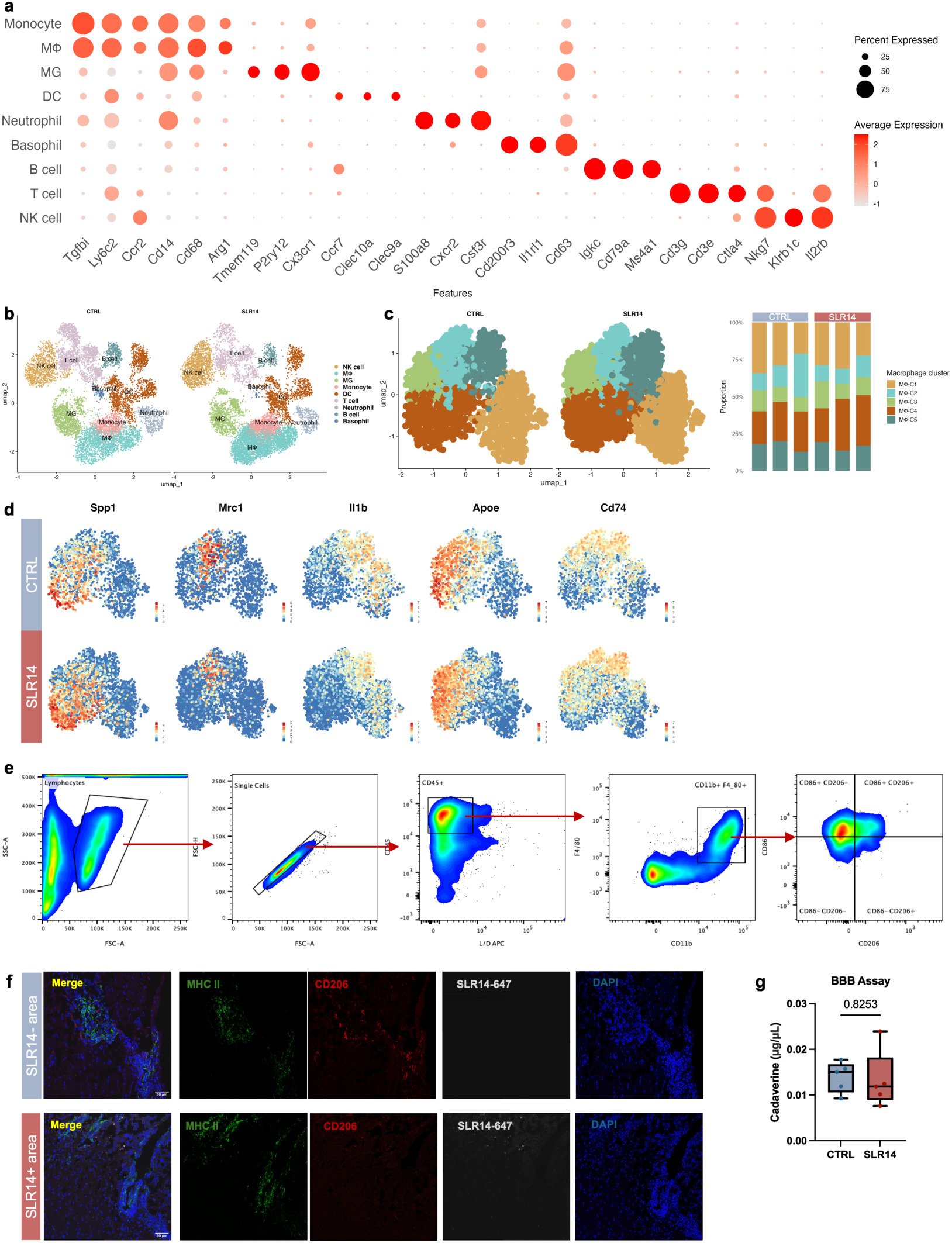
Single-cell RNA-seq analysis of tumor infiltrating immune cells after JetPEI vehicle, or SLR14 treatment. **a**, Gene-expression patterns across individual immune-cell clusters from scRNA-seq of GL261-bearing mice. **b**, UMAP of immune cell from mice treated with vehicle alone or SLR14. **c**. UMAP showing clustering (left) and bar plots showing frequencies (right, scRNA-Seq) of the indicated macrophage types in vehicle (CTRL), or SLR14-treated mice. **d**, UMAP of selected TAMs marker genes’ expression (*Spp1, Mrc1, Il1b, Apoe, Cd74)*, comparing vehicle (CTRL) and SLR14-treated mice. **e**, Gating strategy for M1/M2 macrophage analysis. **f**, Representative immunofluorescence images of MHCII^+^ and CD206^+^ cells within either SLR14-negative or positive areas of the tumor. MHCII (green), CD206 (red), SLR14-647 (gray), and DAPI (blue). Scale bar, 50μm. **g**, Blood-brain barrier (BBB) permeability assay performed on brain section after systemic injection of cadaverine tracer in SLR14-treated mice.

**Extended Data Fig. 6.**
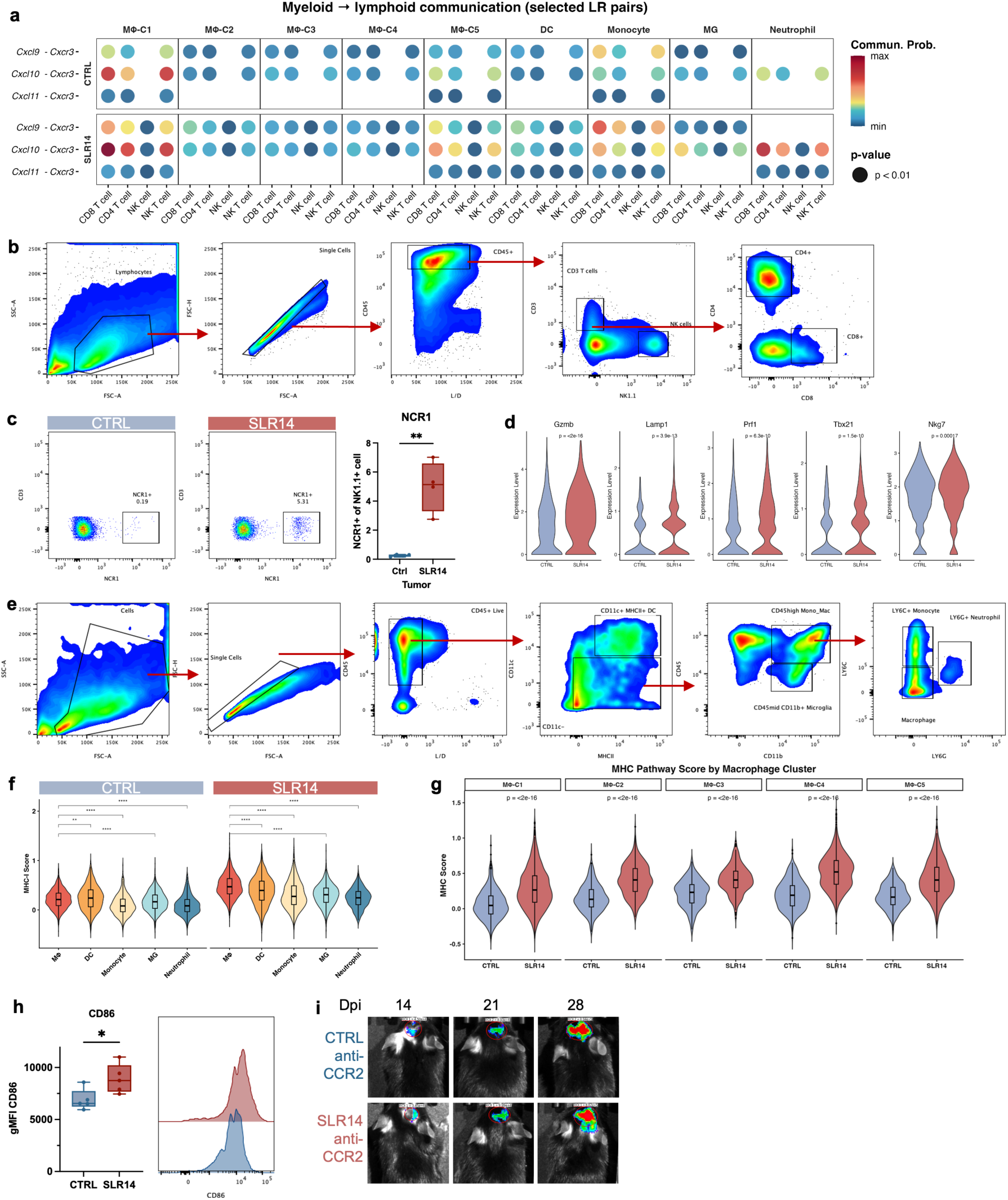
SLR14 enhances antigen presentation and NK cell activation in GBM. **a**, Ligand-receptor pairs contributing to chemokine signaling from macrophage clusters to lymphoid population. The dot color represents communication probability, and size indicates *p* value for interaction significance. **b**, Gate strategy for NK cells, and T cells. **c**, Contour plots (left) and frequency (right) of NK cells (n = 5 per group) and NCR1^+^ NK cells within tumor (n = 4 per group). **d**, Violin plots of NK cell activation genes (*Gzmb, Lamp1, Prf1, Tbx21, Nkg7*) expression in NK cells in SLR14 treated mice compared to control. **e**, Gate strategy for myeloid cells. **f**, Violin plot showing the distribution of MHCI score in different myeloid clusters in each condition. **g**, Violin plot showing the distribution of MHCI score in different macrophage clusters in each condition. **h**, gMFI of CD86 of macrophage and representative histogram from control and SLR14-treated mice (n = 5 per group). **i,** Representative bioluminescence images showing GL261-Luc tumor burden.

**Extended Data Fig. 7.**
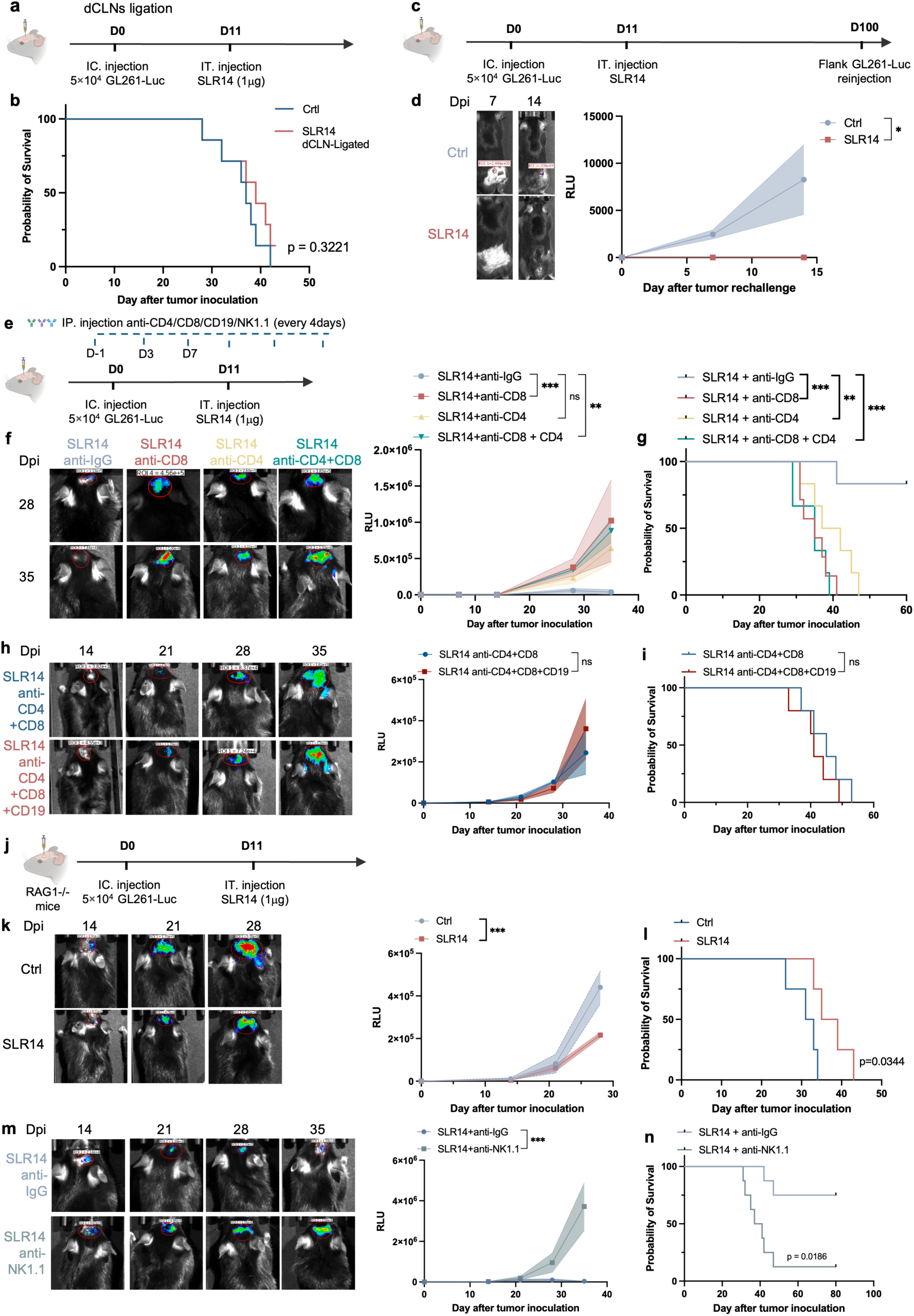
Antitumor effect of SLR14 against GL261-GBM is mainly mediated by T cells. **a**, Schematic for the experimental design in (**b**), using the GL261 GBM model. **b**, Kaplan–Meier survival curves (n = 7 per group). **c**, Schematic for experimental data in (**d**), using the GL261 GBM model. Mice inoculated intracranially with GL261-Luc cells (D0) were treated with vehicle or SLR14 (4 μg) at D11. Surviving mice were rechallenged with GL261 cells by subcutaneous flank injection. **d**, Representative bioluminescence images showing GL261 tumor burden (left) and quantification of tumor volume (right) (n = 5 per group). **e**, Scheme for T cell, B cell or NK cell blockage experiment design in (**f-n**), using the GL261 GBM**. f**, Representative bioluminescence images showing GL261-Luc tumor burden (left) and quantification of tumor volume (right) (n = 6 per group). **g**, Kaplan–Meier survival curves following SLR14 treatment and T cell depletion (n = 6 per group). **h**, Representative bioluminescence images showing GL261-Luc tumor burden (left) and quantification of tumor volume (right) (n = 5 per group). **i**, Kaplan–Meier survival curves of following SLR14 treatment and T cell/B cell depletion (n = 5 per group). **j**, Schematic for experimental data in (**k, l)**. **k**, Representative bioluminescence images showing GL261-Luc tumor burden (left) and quantification of tumor volume (right) (n = 4 per group). **l**, Kaplan–Meier survival curves in RAG1-/- mice following SLR14 treatment (n = 4 per group). **m**, Representative bioluminescence images showing GL261-Luc tumor burden (left) and quantification of tumor volume (right) (n = 4 per group). **n**, Kaplan–Meier survival curves following SLR14 treatment and NK cell depletion (n = 8 per group).

**Extended Data Fig. 8.**
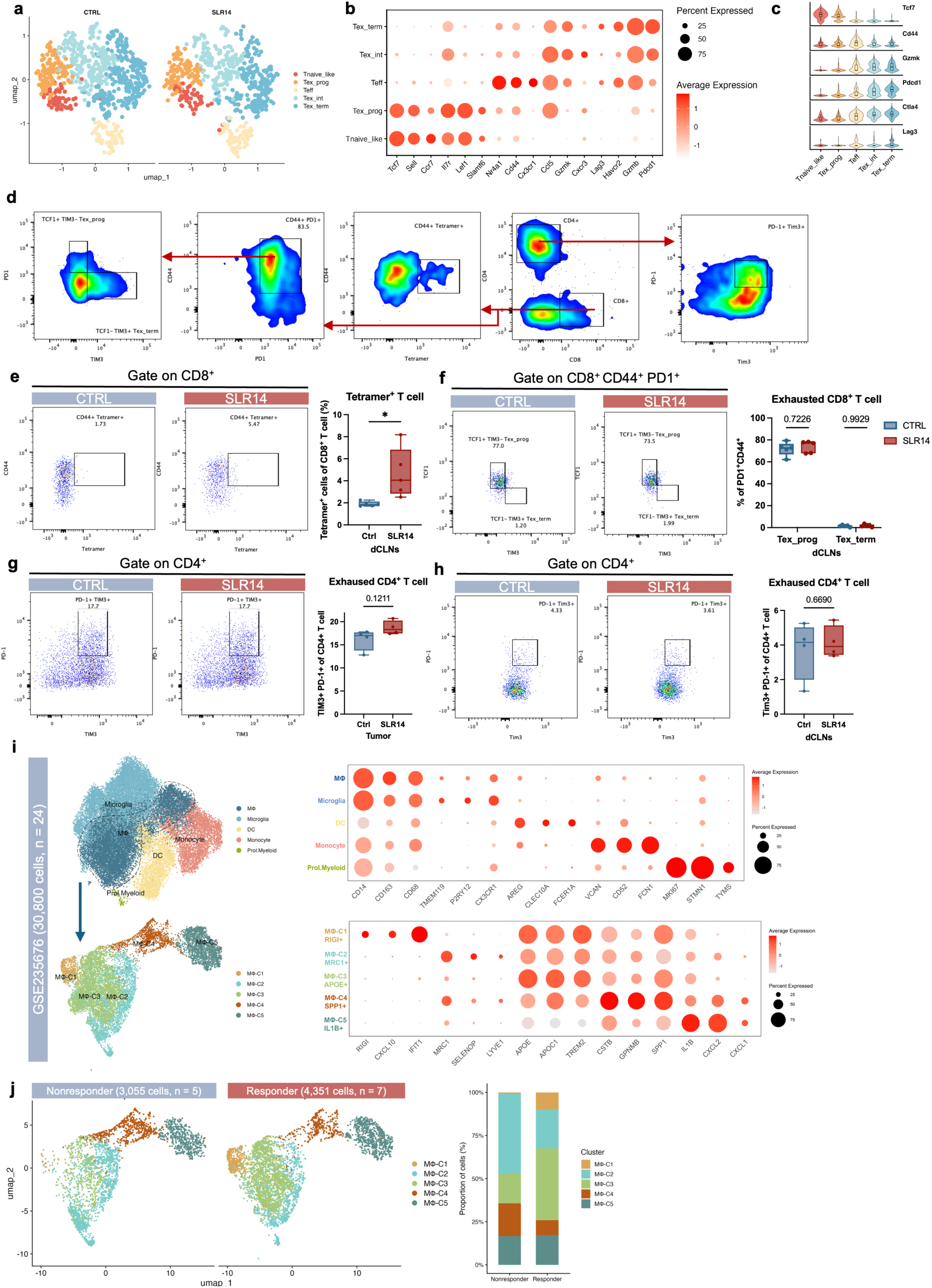
Immune profiling of T cell- and myeloid cell-subsets following anti-PD-1 treatment. **a**, UMAP of CD8 T cells from mice treated with vehicle alone or SLR14. **b**, Gene-expression patterns across CD8^+^ T cell clusters from scRNA-seq of GL261-bearing mice. **c**, Violine plot of *Tcf7, Cd44, Gzmk, Pdcd1, Ctla4,* and *Lag3* expression across clusters. **d**, Gate strategy for functional and exhausted T cells. **e**, Contour plots (left) and frequency (right) of tetramer^+^ T cells in CD8^+^ T cells within dCLNs (n = 5 per group). **f**, Contour plots (left), and frequency (right) of TCF1^+^TIM3^+^ Tex-prog and TCF1^+^TIM3^+^ Tex-term cells in CD8^+^ CD44^+^ PD1^+^ T cells within dCLNs (n = 5 per group). **g**, Contour plots (left) and frequency (right) of exhausted CD4^+^ T cells within tumor (n = 4 per group). **h**, Contour plots (left) and frequency (right) of exhausted CD4^+^ T cells within dCLNs (n = 4 per group). **i**, Clustering of myeloid cell types and TAM clusters (left) and gene-expression patterns (right) using public scRNA-seq datasets of GBM in human. **j**, UMAP showing clustering (left) and bar plots showing frequencies (right, scRNA-Seq) of indicated macrophage subsets in anti-PD-1-treated responder and nonresponder groups of patients with GBM (GSE235676-

## Reference

1. Price, M., et al. CBTRUS Statistical Report: Primary Brain and Other Central Nervous System Tumors Diagnosed in the United States in 2018-2022. Neuro Oncol 27, iv1–iv66 (2025). 10.1093/neuonc/noaf194

2 Reardon, D. A. et al. Effect of Nivolumab vs Bevacizumab in Patients With Recurrent Glioblastoma: The CheckMate 143 Phase 3 Randomized Clinical Trial. JAMA Oncol 6, 1003–1010 (2020). 10.1001/jamaoncol.2020.1024

3 Preusser, M., Lim, M., Hafler, D. A., Reardon, D. A. & Sampson, J. H. Prospects of immune checkpoint modulators in the treatment of glioblastoma. Nat Rev Neurol 11, 504–514 (2015). 10.1038/nrneurol.2015.139

4 Ribas, A. & Wolchok, J. D. Cancer immunotherapy using checkpoint blockade. Science 359, 1350–1355 (2018). 10.1126/science.aar4060

5 Chinot, O. L. et al. Bevacizumab plus radiotherapy-temozolomide for newly diagnosed glioblastoma. N Engl J Med 370, 709–722 (2014). 10.1056/NEJMoa1308345

6 Grossman, S. A. et al. Immunosuppression in patients with high-grade gliomas treated with radiation and temozolomide. Clin Cancer Res 17, 5473–5480 (2011). 10.1158/1078-0432.Ccr-11-0774

7 Wang, A. Z. et al. Glioblastoma-Infiltrating CD8+ T Cells Are Predominantly a Clonally Expanded GZMK+ Effector Population. Cancer Discovery 14, 1106–1131 (2024). 10.1158/2159-8290.Cd-23-0913

8 Wang, W. et al. Identification of hypoxic macrophages in glioblastoma with therapeutic potential for vasculature normalization. Cancer Cell 42, 815–832.e812 (2024). 10.1016/j.ccell.2024.03.013

9 Galvez-Cancino, F. et al. Regulatory T cell depletion promotes myeloid cell activation and glioblastoma response to anti-PD1 and tumor-targeting antibodies. Immunity 58, 1236–1253.e1238 (2025). 10.1016/j.immuni.2025.03.021

10 Khasraw, M., Reardon, D. A., Weller, M. & Sampson, J. H. PD-1 Inhibitors: Do they have a Future in the Treatment of Glioblastoma? Clinical Cancer Research 26, 5287–5296 (2020). 10.1158/1078-0432.Ccr-20-1135

11 Woroniecka, K. et al. T-Cell Exhaustion Signatures Vary with Tumor Type and Are Severe in Glioblastoma. Clin Cancer Res 24, 4175–4186 (2018). 10.1158/1078-0432.Ccr-17-1846

12 Medzhitov, R. The spectrum of inflammatory responses. Science 374, 1070–1075 (2021). 10.1126/science.abi5200

13 Caronni, N. et al. IL-1β+ macrophages fuel pathogenic inflammation in pancreatic cancer. Nature 623, 415–422 (2023). 10.1038/s41586-023-06685-2

14 Li, D. & Wu, M. Pattern recognition receptors in health and diseases. Signal Transduction and Targeted Therapy 6, 291 (2021). 10.1038/s41392-021-00687-0

15 Rehwinkel, J. & Gack, M. U. RIG-I-like receptors: their regulation and roles in RNA sensing. Nat Rev Immunol 20, 537–551 (2020). 10.1038/s41577-020-0288-3

16 Demaria, O. et al. Harnessing innate immunity in cancer therapy. Nature 574, 45–56 (2019). 10.1038/s41586-019-1593-5

17 Berger, G. et al. STING activation promotes robust immune response and NK cell-mediated tumor regression in glioblastoma models. Proc Natl Acad Sci U S A 119, e2111003119 (2022). 10.1073/pnas.2111003119

18 Najem, H. et al. STING agonist 8803 reprograms the immune microenvironment and increases survival in preclinical models of glioblastoma. The Journal of Clinical Investigation 134 (2024). 10.1172/JCI175033

19 Sun, S. et al. Targeting GOLPH3L improves glioblastoma radiotherapy by regulating STING-NLRP3-mediated tumor immune microenvironment reprogramming. Sci Transl Med 17, eado0020 (2025). 10.1126/scitranslmed.ado0020

20 Everson, R. G. et al. TLR agonists polarize interferon responses in conjunction with dendritic cell vaccination in malignant glioma: a randomized phase II Trial. Nature Communications 15, 3882 (2024). 10.1038/s41467-024-48073-y

21 Elion, D. L. et al. Therapeutically Active RIG-I Agonist Induces Immunogenic Tumor Cell Killing in Breast Cancers. Cancer Research 78, 6183–6195 (2018). 10.1158/0008-5472.Can-18-0730

22 Hou, J. et al. Hepatic RIG-I Predicts Survival and Interferon-α Therapeutic Response in Hepatocellular Carcinoma. Cancer Cell 25, 49–63 (2014). 10.1016/j.ccr.2013.11.011

23 Chen, M. et al. Targeting nuclear acid-mediated immunity in cancer immune checkpoint inhibitor therapies. Signal Transduction and Targeted Therapy 5, 270 (2020). 10.1038/s41392-020-00347-9

24 Linehan, M. M. et al. A minimal RNA ligand for potent RIG-I activation in living mice. Science Advances 4, e1701854 (2018). doi:10.1126/sciadv.1701854

25 Jiang, X. et al. Intratumoral delivery of RIG-I agonist SLR14 induces robust antitumor responses. Journal of Experimental Medicine 216, 2854–2868 (2019). 10.1084/jem.20190801

26 Abdelfattah, N. et al. Single-cell analysis of human glioma and immune cells identifies S100A4 as an immunotherapy target. Nature Communications 13, 767 (2022). 10.1038/s41467-022-28372-y

27 Siret, C. et al. Deciphering the heterogeneity of the Lyve1+ perivascular macrophages in the mouse brain. Nature Communications 13, 7366 (2022). 10.1038/s41467-022-35166-9

28 Bijnen, M., Sridhar, S., Keller, A. & Greter, M. Brain macrophages in vascular health and dysfunction. Trends in Immunology 46, 46–60 (2025). 10.1016/j.it.2024.11.012

29 Bill, R. et al. CXCL9:SPP1 macrophage polarity identifies a network of cellular programs that control human cancers. Science 381, 515–524 (2023). 10.1126/science.ade2292

30 Ravi, V. M. et al. Spatially resolved multi-omics deciphers bidirectional tumor-host interdependence in glioblastoma. Cancer Cell 40, 639–655.e613 (2022). 10.1016/j.ccell.2022.05.009

31 Van Hove, H. et al. Interleukin-34-dependent perivascular macrophages promote vascular function in the brain. Immunity 58, 1289–1305.e1288 (2025). 10.1016/j.immuni.2025.04.003

32 Onomoto, K., Onoguchi, K. & Yoneyama, M. Regulation of RIG-I-like receptor-mediated signaling: interaction between host and viral factors. Cell Mol Immunol 18, 539–555 (2021). 10.1038/s41423-020-00602-7

33 Jiang, Y. et al. Exploiting RIG-I-like receptor pathway for cancer immunotherapy. J Hematol Oncol 16, 8 (2023). 10.1186/s13045-023-01405-9

34 Genoud, V. et al. Responsiveness to anti-PD-1 and anti-CTLA-4 immune checkpoint blockade in SB28 and GL261 mouse glioma models. Oncoimmunology 7, e1501137 (2018). 10.1080/2162402x.2018.1501137

35 Jin, S., Plikus, M. V. & Nie, Q. CellChat for systematic analysis of cell–cell communication from single-cell transcriptomics. Nature Protocols 20, 180–219 (2025). 10.1038/s41596-024-01045-4

36 Song, E. et al. VEGF-C-driven lymphatic drainage enables immunosurveillance of brain tumours. Nature 577, 689–694 (2020). 10.1038/s41586-019-1912-x

37 Hu, X. et al. Meningeal lymphatic vessels regulate brain tumor drainage and immunity. Cell Res 30, 229–243 (2020). 10.1038/s41422-020-0287-8

38 Zhou, C., Ma, L., Xu, H., Huo, Y. & Luo, J. Meningeal lymphatics regulate radiotherapy efficacy through modulating anti-tumor immunity. Cell Res 32, 543–554 (2022). 10.1038/s41422-022-00639-5

39 Bronte, V. et al. Effective genetic vaccination with a widely shared endogenous retroviral tumor antigen requires CD40 stimulation during tumor rejection phase. J Immunol 171, 6396–6405 (2003). 10.4049/jimmunol.171.12.6396

40 Mei, Y. et al. Siglec-9 acts as an immune-checkpoint molecule on macrophages in glioblastoma, restricting T-cell priming and immunotherapy response. Nature Cancer 4, 1273–1291 (2023). 10.1038/s43018-023-00598-9

41 Quail, D. F. & Joyce, J. A. The Microenvironmental Landscape of Brain Tumors. Cancer Cell 31, 326–341 (2017). 10.1016/j.ccell.2017.02.009

42 Waibl Polania, J., et al. Antigen presentation by tumor-associated macrophages drives T cells from a progenitor exhaustion state to terminal exhaustion. Immunity 58, 232–246.e236 (2025). 10.1016/j.immuni.2024.11.026

43 Lee, A. H. et al. Neoadjuvant PD-1 blockade induces T cell and cDC1 activation but fails to overcome the immunosuppressive tumor associated macrophages in recurrent glioblastoma. Nat Commun 12, 6938 (2021). 10.1038/s41467-021-26940-2

44 Rodriguez-Baena, F. J. et al. Microglial reprogramming enhances antitumor immunity and immunotherapy response in melanoma brain metastases. Cancer Cell 43, 413–427.e419 (2025). 10.1016/j.ccell.2025.01.008

45 Cao, J. et al. The single-cell transcriptional landscape of mammalian organogenesis. Nature 566, 496–502 (2019). 10.1038/s41586-019-0969-x

46 Greenwald, A. C. et al. Integrative spatial analysis reveals a multi-layered organization of glioblastoma. Cell 187, 2485–2501.e2426 (2024). 10.1016/j.cell.2024.03.029

